# Epigenetic modulation of NLRP6 inflammasome sensor as a therapeutic modality to reduce necroptosis-driven gastrointestinal mucosal dysfunction in HIV/SIV infection

**DOI:** 10.1101/2024.11.13.623322

**Authors:** Lakmini S. Premadasa, Marina McDew-White, Luis Romero, Beverly Gondo, Jade A. Drawec, Binhua Ling, Chioma M. Okeoma, Mahesh Mohan

## Abstract

The epigenetic mechanisms driving persistent gastrointestinal mucosal dysfunction in HIV/SIV infection is an understudied topic. Using reduced-representation bisulfite sequencing, we identified HIV/SIV infection in combination anti-retroviral therapy (cART)-naive rhesus macaques (RMs) to induce marked hypomethylation throughout promoter-associated CpG islands (paCGIs) in genes related to inflammatory response (*NLRP6, cGAS*), cellular adhesion and proliferation in colonic epithelial cells (CEs). Moreover, low-dose delta-9-tetrahydrocannabinol (THC) administration reduced NLRP6 protein expression in CE by hypermethylating the *NLRP6* paCGI and blocked polyI:C induced NLRP6 upregulation in vitro. In cART suppressed SIV-infected RMs, NLRP6 protein upregulation associated with significantly increased expression of necroptosis-driving proteins; phosphorylated-RIPK3(Ser199), phosphorylated-MLKL(Thr357/Ser358), and HMGB1. Most strikingly, supplementing cART with THC effectively reduced NLRP6 and necroptosis-driving protein expression to pre-infection levels. These findings for the first time demonstrate that NLRP6 upregulation and ensuing activation of necroptosis promote HIV/SIV-induced gastrointestinal mucosal dysfunction and that epigenetic modulation using phytocannabinoids represents a feasible therapeutic modality for alleviating HIV/SIV-induced gastrointestinal inflammation and associated comorbidities.

## Introduction

The gastrointestinal (GI) tract, the predominant site of human/simian immunodeficiency virus (HIV/SIV) replication, persistence, and dissemination, plays the triple role of nutrient, water, and anti-retroviral drug absorption while simultaneously protecting the host from pathogenic bacteria and their metabolites. Chronic inflammation of the colon, home to trillions of commensal bacteria, can perturb the homeostatic balance between the host and gut microbiota, causing dysbiosis and epithelial barrier dysfunction that can lead to microbial translocation, which can perpetuate systemic inflammation ^1^. Despite viral suppression by combination anti-retroviral therapy (cART), HIV/SIV-associated enteropathy that persists in people living with HIV (PLWH)^2^ may result in systemic inflammation that can contribute to numerous other non-AIDS-associated comorbidities, thereby decreasing their overall quality of life ^1,3–5^. Persistent GI inflammation has also been shown to occur in people living with inflammatory bowel disease (IBD), despite the absence of overt GI symptoms ^6^. More importantly, inflammation-driven chronic intestinal barrier impairment has been linked to extra-intestinal comorbidities, such as nonalcoholic fatty liver disease and type-2 diabetes, and is associated with neurocognitive and inflammatory diseases^1^ including Parkinson’s disease, Alzheimer’s disease, and multiple sclerosis ^7^. We previously showed that an impaired intestinal barrier caused by HIV/SIV-induced intestinal inflammation can alter signaling along the microbiota-gut-brain-axis, where dysbiotic microbes and microbial-derived by-products can increase intestinal inflammation, increase type-I interferon responses, endoplasmic reticulum, and oxidative stress in the neurons of the basal ganglia, and could lead to neuroinflammation and potentially cognitive decline in PLWH ^8^.

While the molecular mechanisms underlying GI epithelial dysfunction in PLWH are ill-defined, emerging evidence in other GI inflammatory diseases, such as IBD and irritable bowel syndrome (IBS), points to epigenetic origins. In this context, aberrant DNA methylation and non-coding RNAs (microRNAs and long non-coding RNAs) have been shown to contribute to epithelial dysfunction in IBD and irritable bowel syndrome (IBS) ^9–31^. Moreover, therapeutic drugs to treat intestinal epithelial barrier impairment, and restore barrier function are currently unavailable, and therefore, this topic is of significant interest for future drug development ^6,7^. Due to the heritable nature of epigenetic changes, the role of DNA methylation in chronic intestinal inflammation needs to be carefully examined, and promising immune modulation strategies should be explored to reduce inflammation and disease progression in not only PLWH but also those who suffer from IBD, as only 40-60% of the population respond to current therapies ^32^.

Here, we used reduced representation bisulfite sequencing (RRBS) to better understand how epigenetic regulation, through DNA methylation of colonic epithelial genes, can impact pro-inflammatory genes and lead to chronic intestinal inflammation and epithelial dysfunction in SIV-infected rhesus macaques (RMs). While extremely challenging in PLWH, intestinal resections (∼5 cm) were collected from SIV-infected RMs without any adverse effects, providing a unique avenue to identify epigenetically altered protein-coding genes involved in intestinal barrier disruption ^33–39^. We found marked alterations in DNA methylation, specifically hypomethylation occurring in promoter-associated CpG islands (paCGI), in SIV-infected RMs. These genes were linked to epithelial cell proliferation and adhesion, apoptosis, inflammatory response, dsDNA damage response, and oxidative stress. More importantly, we showed that long-term, low-dose phytocannabinoid administration to SIV-infected RMs successfully modulated methylation in paCGIs, notably hypermethylation in the CpG island of the NOD-like receptor family pyrin domain containing 6 (NLRP6), which markedly decreased protein expression of NLRP6, a sensor component of the NLRP6 inflammasome known to maintain intestinal epithelial barrier function and protect against microbial invasion, relative to control RMs. Persistent NLRP6 activation was associated with significantly high mRNA and protein expression of necroptosis-promoting RIPK1-RIPK3-MLKL pathway. In in vitro cultured human small intestinal epithelial cells, both THC and cannabidiol (CBD) successfully blocked NLRP6 protein upregulation in response to polyI:C and lipoteichoic acid. Lastly, while long-term cART administration alone did not fully reduce NLRP6 protein expression in the intestines of SIV-infected RMs, supplementing long-term cART with low-dose THC successfully reduced NLRP6 protein compared to the increase seen one month after infection.

Our findings provide novel and translational insights into the epigenetic regulation of intestinal epithelial gene expression in HIV/SIV infection and phytocannabinoid-induced alteration of DNA methylation as an important underreported molecular mechanism attributable to its anti-inflammatory and cellular protective properties. Benefits could also expand to other diseases associated with intestinal inflammation, such as IBD, where interventions to address the cause and restoration of barrier integrity have not been successful. Additionally, we show that chronic low-dose phytocannabinoid administration could be used as a safe ^40^ intervention and restorative therapy for persistent intestinal inflammation, and its use alone or as an adjunct to cART can be beneficial for PLWH^40^ with no access to cART or to those who fail to fully suppress HIV under cART.

## Results

### Plasma, colon, and jejunum viral loads and histopathology

cART-naïve RMs in groups 2 and 3 had substantial plasma (0.01 x 10^6^ to 2.6 x 10^8^) and colon (1 x 10^6^ to 9.36 x 10^9^) viral loads at 6MPI. cART-treated RMs in groups 4 and 5 had undetectable plasma and jejunum viral loads at 5MPI (Table 1). Histopathological analysis revealed moderate colitis and cryptitis and lymphoid hyperplasia in four group 2 and two group 3 RMs (Table 1). SIV syncytial cells were detected in one group 3 macaque. No opportunistic infections were detected in any animals.

**Table 1.** Animal IDs, SIV infection (SIVmac251) duration time, cannabinoids and cART treatment, viral load for plasma, colon, and jejunum, and histopathology for colon and jejunum tissue.

| Animal ID | Duration of Infection | Plasma viral loads 10<sup>6</sup>/mL | Colon viral loads 10<sup>6</sup>/mg RNA | Jejunum viral loads 10<sup>6</sup>/mg RNA | Colon Histopathology | Jejunum Histopathology |
| --- | --- | --- | --- | --- | --- | --- |
| **Group 1: Uninfected Controls (Controls)** |  |  |  |  |  |  |
| CONT1, CONT2, CONT3, CONT4<sup>a</sup> | NA | NA | NA | NA | NA | NA |
| CONT5<sup>a,b</sup> | NA | NA | NA | NA | NA | NA |
| CONT6<sup>b</sup> | NA | NA | NA | NA | NA | NA |
| CONT7<sup>*b</sup> | NA | NA | NA | NA | NA | NA |
| CONT8<sup>*b</sup> | NA | NA | NA | NA | NA | NA |
| CONT9, CONT10, CONT11<sup>c</sup> | NA | NA | NA | NA | NA | NA |
| CONT12, CONT13, CONT14<sup>c</sup> | NA | NA | NA | NA | NA | NA |
| CONT15, CONT16<sup>c</sup> | NA | NA | NA | NA | NA | NA |
| **Group 2: Chronic SIV-infected and Vehicle Treated (VEH/SIV)** |  |  |  |  |  |  |
| VEHSIV1<sup>a,c</sup> | 150 | 4 | 147 | NA | ND | NA |
| VEHSIV2<sup>a</sup> | 180 | 0.02 | 2 | NA | ND | NA |
| VEHSIV3<sup>a</sup> | 180 | 2 | 300 | NA | ND | NA |
| VEHSIV4<sup>b,c</sup> | 180 | 0.1 | 786 | NA | Lymphoid hyperplasia | NA |
| VEHSIV5<sup>a,b,c</sup> | 180 | 9.4 | 29 | NA | ND | NA |
| VEHSIV6<sup>a,b,c</sup> | 180 | 0.38 | 320 | NA | ND | NA |
| VEHSIV7<sup>a,c</sup> | 180 | 0.04 | 4 | NA | ND | NA |
| VEHSIV8<sup>a,c</sup> | 180 | 0.5 | 20 | NA | ND | NA |
| VEHSIV9<sup>*b</sup> | 90 | 37 | 3000 | NA | Moderate colitis/cryptitis | NA |
| VEHSIV10<sup>*b,c</sup> | 180 | 9 | 59 | NA | ND | NA |
| VEHSIV11<sup>c</sup> | 180 | 30 | 3 | NA | Lymphoid hyperplasia | NA |
| VEHSIV12<sup>c</sup> | 180 | 3000 | 200 | NA | Lymphoid hyperplasia | NA |
| VEHSIV13<sup>c</sup> | 180 | 0.2 | 210 | NA | ND | NA |
| VEHSIV14<sup>b</sup> | 180 | NA | 11 | NA | ND | NA |
| **Group 3: Chronic SIV-infected and THC treated (THC/SIV)** |  |  |  |  |  |  |
| THCSIV1<sup>b</sup> | 180 | 18.9 | 6726 | NA | Mild hyperplasia/SIV syncytia | NA |
| THCSIV2<sup>a</sup> | 150 | 260 | 9,360 | NA | ND | NA |
| THCSIV3<sup>a</sup> | 180 | 3 | 10 | NA | ND | NA |
| THCSIV4<sup>a,b,c</sup> | 180 | 0.01 | 3 | NA | ND | NA |
| THCSIV5<sup>b,c</sup> | 180 | 0.06 | 93.8 | NA | ND | NA |
| THCSIV6<sup>a,c</sup> | 180 | 0.02 | 10 | NA | ND | NA |
| THCSIV7<sup>a,c</sup> | 180 | 0.02 | 1 | NA | ND | NA |
| THCSIV8<sup>a,c</sup> | 180 | 1 | 300 | NA | ND | NA |
| THCSIV9<sup>b</sup> | 180 | 18.9 | 6,726 | NA | ND | NA |
| THCSIV10<sup>b,c</sup> | 180 | 1.5 | 1,261 | NA | ND | NA |
| THCSIV11<sup>b,c</sup> | 180 | 7.7 | 970 | NA | ND | NA |
| THCSIV12<sup>c</sup> | 150 | 0.66 | 35 | NA | Mild colitis | NA |
| **Group 4: Chronic SIV-infected, cART and Vehicle treated (VEH/SIV/cART)** |  |  |  |  |  |  |
| VEHSIVART1, VEHSIVART2,<br>VEHSIVART3, VEHSIVART4,<br>VEHSIVART5, VEHSIVART6,<br>VEHSIVART7, VEHSIVART8<sup>*c</sup> | 150 | ND | NA | ND | NA | ND |
| **Group 5: Chronic SIV-infected and THC treated and cART (THC/SIV/cART)** |  |  |  |  |  |  |
| THCSIVART1, THCSIVART2,<br>THCSIVART3, THCSIVART4,<br>THCSIVART5, THCSIVART6,<br>THCSIVART7, THCSIVART8<sup>*c</sup> | 150 | ND | NA | ND | NA | ND |

NA- not applicable, ND- none detected
\* denotes animals pre-infection timepoint used as control
<sup>a</sup> denotes animals used for reduced representation bisulfite sequencing study
<sup>b</sup> denotes animals used for microarray gene expression study
<sup>c</sup> denotes animals used for NLRP6 and necroptosis immunofluorescence (confocal) studies

### Chronic HIV/SIV infection is characterized by global DNA hypomethylation concentrated mainly in promoter-associated CpG islands in colonic epithelium

Based on findings from DNA methylation profiling of CE using RRBS, relative to controls, both VEH/SIV and THC/SIV RMs, and VEH/SIV compared to THC/SIV RMs showed more changes in CpG methylation ratios [Δβ-value] (a decrease indicates hypomethylation change, while an increase indicates hypermethylation change); p-value <0.05, absolute Δβ-value >0.1) distributed throughout all chromosomes, with the least amount of differentially methylated CpG sites (DMSs) occurring in chromosome Y (Figure 1A). All data presented below is described in the flow chart in Figure S1. As shown in Figure 1A, compared to controls, THC/SIV RMs had more DMSs than VEH/SIV RMs. In addition, THC/SIV RMs showed more hypomethylation (values below 0) changes in all comparisons than hypermethylation (values above 0) (Figure 1A). Only DMSs within annotated genes (intergenic and unannotated regions excluded) were analyzed. Compared to controls, VEH/SIV RMs had a total of 13,831 [out of 94,279 (14.67%)] DMSs within annotated genes (Figure 1B). A similar number of DMSs were detected in THC/SIV compared to control RMs; 13.59% of DMSs were within annotated genes (16,730 out of 123,135 DMSs; Figure 1C). Lastly, compared to THC/SIV, VEH/SIV RMs had a total of 6,796 [out of 64,069 (10.61%)] DMSs within annotated genes (Figure 1D).

**Figure 1.**
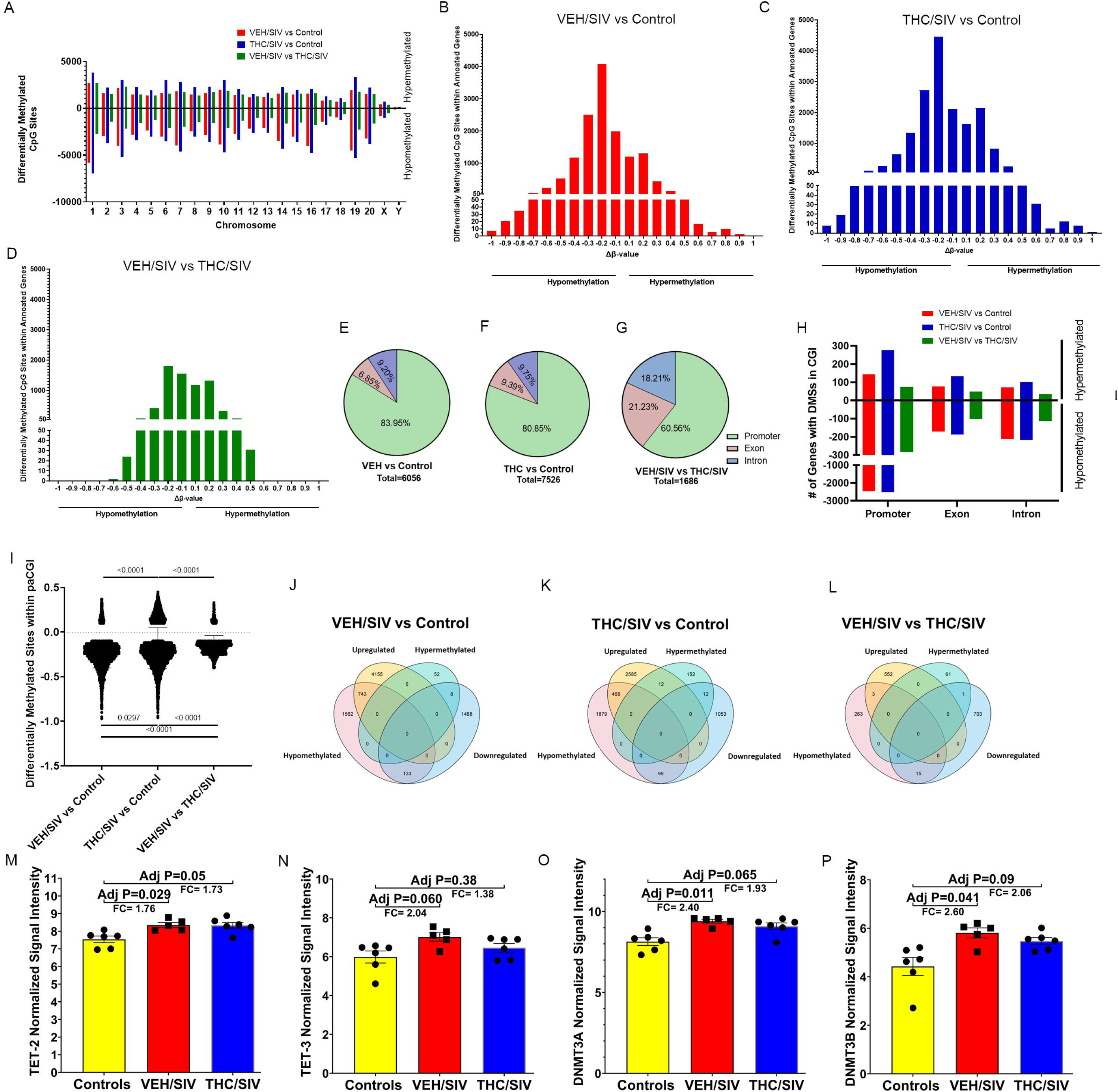
Hypomethylation changes in promoter associated CpG islands (paCGIs) associated with upregulation of their respective gene and methylation altering machinery gene expression. (**A**) Distribution of differentially methylated sites (DMSs) (p-value <0.05, absolute Δβ-value >0.1) in individual chromosomes of the rhesus macaque (RM) genome. After excluding intergenic DMSs, the bar graphs show the remaining 14.67% (**B**), 13.59% (**C**) and 10.61% (**D**) DMSs in annotated genes of VEH/SIV (**B**; red) and THC/SIV (**C**; blue) compared to control or VEH/SIV compared to THC/SIV (**D**; gold) RMs, respectively. Percentage of DMSs present in the promoter, exon and intronic regions contained within CpG islands (CGI) of annotated genes in VEH/SIV (**E)** and THC/SIV (**F**) compared to control or VEH/SIV (**G**) compared to THC/SIV RMs. Number of genes that showed hypo or hypermethylation in their paCGIs compared to their respective exons and introns in VEH/SIV (red) and THC/SIV (blue) relative to control or VEH/SIV compared to THC/SIV RMs (gold) (**H**). Δβ-value of DMSs within the paCGIs between VEH/SIV and THC/SIV compared to controls, and THC/SIV relative to control vs VEH/SIV compared to THC/SIV RMs. Venn diagram showing relationship between Δβ-values of DMSs in paCGIs and gene expression in VEH/SIV (**J**) and THC/SIV (**K**) relative to control or VEH/SIV compared to THC/SIV RMs (**L**). Normalized signal intensity and fold change of *TET2* (**M**), *TET3* (**N**), *DNMT3A* (**O**) and *DNMT3B* (**P**) mRNA in colonic epithelium of VEH/SIV and THC/SIV compared to control RMs. Differences in methylation changes between groups were analyzed using Kruskal-Wallis test (GraphPad Prism). A p-value of <0.05 was considered significant. Data represented as mean ± SD.

We focused on methylation changes that occurred within the CpG-rich genomic regions called CpG islands (CGIs) that also included the promoter, exon, and intron intragenic regions. CGIs that spanned a promoter region (continuing into the exon and intron regions) were classified as paCGIs. Nearly 70% of gene promoters reside within CGIs and are normally unmethylated ^41–43^. Interestingly, paCGIs are devoid of common promoter elements (TATA boxes and core promoter elements), but contain binding sites for transcription factors, which help their respective genes to maintain a transcriptionally permissive state. Out of all the DMSs within annotated genes, VEH/SIV RMs had 43.79% located within CGIs (6,056 out of 13,831), relative to control RMs. These DMSs were identified in 2,688 genes, of which 2,504 genes had 5,084 (83.95%) DMSs within paCGIs (Figure 1E and 1H). Relative to controls, THC/SIV RMs showed 7,526 out of 16,730 (44.99%) DMSs in CGIs within annotated genes. There were 2,622 unique genes that had 80.85% (6,085 out of 7,526) of DMSs in paCGIs within annotated genes (Figure 1F and 1H). Finally, VEH/SIV RMs had 1,686 (24.81%) DMSs within 540 CGIs of annotated genes and contained 1,021 (60.56%) DMSs within the paCGIs of 343 genes, relative to THC/SIV RMs (Figure 1G and 1H).

Intriguingly, untreated HIV/SIV infection was associated with more hypomethylation changes in the paCGI [Figure 1B, 1C (Δβ-values to the left of 0 on X-axis), 1H, and 1I (Δβ-values below 0)]. The largest Δβ-value change detected was a decrease of 20% (Figure 1B and 1C), and both VEH/SIV and THC/SIV RMs had more promoters that were hypomethylated than exons and introns in CGIs (Figure 1H and 1I). Relative to controls, THC/SIV RMs also showed increased hypermethylation in the promoters, exons, and introns than VEH/SIV RMs (Figure 1H). Interestingly, all comparisons showed a large range of hypomethylation changes that occurred in the paCGI (10-90% Δβ-value decrease), but hypermethylation changes were restricted to less than 50% (10-45% Δβ-value increase, Figure 1I). There was also a significant difference in the total number and hypomethylation changes detected in the paCGIs of VEH/SIV and THC/SIV RMs compared to controls and in VEH/SIV relative to THC/SIV RMs (Figure 1I). The same was true for the total number and hypermethylation changes that occurred in the paCGIs in VEH/SIV and THC/SIV compared to controls, and in VEH/SIV vs. THC/SIV RMs, relative to THC/SIV compared to controls (Figure 1I).

Figure S2A-C shows clustering heatmaps of the top 100 differentially methylated CpG sites, including those located in intergenic regions, demonstrating major differences in methylation among the three different groups. The top 100 DMSs are mostly hypomethylated in VEH/SIV and THC/SIV relative to control RMs (Figures S2A and S2B). When compared to THC/SIV, the top 100 DMSs in VEH/SIV RMs showed a range of methylation changes (Figure S2C), with hypomethylation predominating. The Pearson’s correlation coefficients for all CpG sites between VEH/SIV vs. control, THC/SIV vs. control, and VEH/SIV vs. THC/SIV RMs were 0.9400, 0.9358, and 0.9359, respectively (Figure S2D-F). These values look very strong, suggesting that each point where the methylation value is different indicates a major treatment effect.

### Hypomethylation of paCGI sites resulted in upregulation of their corresponding gene expression, including the machinery responsible for demethylation and de novo methylation

Next, we assessed the effect of the cumulative Δβ-value (average of all DMSs Δβ-values per gene) of DMSs in individual paCGIs on gene expression. We compared the number of genes that had a cumulative Δβ-value (hypo- or hypermethylation) in the paCGIs to the number of genes that were differentially expressed in our recently published manuscript ^44^ that utilized a human microarray platform designed to profile mRNAs and lncRNAs in the same RMs using CE total RNA. Methylation changes in the paCGIs of genes showed an inverse association with gene regulation (i.e., hypomethylated paCGIs and respective gene upregulation or hypermethylation paCGIs and respective gene downregulation) in 11.5% [751 out of 6533 differentially expressed genes (DEGs)] and 11.4% (480 out of 4229 DEGs) of genes in VEH/SIV and THC/SIV RMs, respectively, compared to controls (Figure 1J and 1K). Hypomethylation was better associated with gene expression changes (increased expression) than hypermethylation, as 99% (743/751 genes were hypomethylated and upregulated, compared to 8/751 genes that were hypermethylated and downregulated) (Figure 1J) and 97.5% (468/480 genes were hypomethylated and upregulated, compared to 12/480 genes that were hypermethylated and downregulated) (Figure 1K) of all genes with DMSs in the paCGIs were hypomethylated in VEH/SIV and THC/SIV RMs, respectively, compared to controls. Surprisingly, methylation changes in the paCGIs had minimal influence on gene expression in VEH/SIV when compared to THC/SIV RMs, as only 4 genes with methylation changes in the paCGIs inversely impacted gene expression (Figure 1L). We also found that hypermethylation in the paCGIs was associated with downregulation of gene expression, as well as an association between hypomethylation and upregulation of gene expression. Consistent with these findings, mRNA upregulation of the enzymes involved in demethylation (TET-2; 1.76-fold) (Figure 1M) and *de novo* methylation [DNMT3a; 2.4-fold (Figure 1O) and DNMT3b; 2.06-fold) (Figure 1P)] was detected in VEH/SIV (n=5) but not THC/SIV (n=6) compared to control RMs (n=6). Although upregulated, TET-3 mRNA did not show statistical significance (Figure 1N).

### Long-term, low-dose THC administration modulated gene expression associated with intercellular signaling, communication, junction organization, and NLRP6 inflammasome assembly

Using Gene Ontology (GO) enrichment analysis, we investigated the effect of the cumulative Δβ-value in the paCGIs of protein-coding genes on the biological processes in CE during HIV/SIV infection. HIV/SIV infection-induced hypomethylation affected the expression of genes linked to epithelial cell proliferation and its regulation, cell growth, cellular response to transforming growth factor-β (TGF-β), regulation of and/or response to oxidative stress, apoptotic signaling pathway, programmed cell death, canonical Wnt signaling pathway, cell communication and signaling, and MAPK cascade (Figures 2A and 2B). Relative to controls, phytocannabinoid administration had little impact on the number of genes hypomethylated in the above biological processes, compared to VEH/SIV RMs (Figures 2A and 2B). When compared to THC/SIV, VEH/SIV RMs had significantly more hypomethylated genes involved in positive regulation of cell communication (Figure 2B). These animals also had hypermethylated genes involved in the overall and negative regulation of cell communication, regulation of cell signaling, and MAPK cascade (Figure 2B).

**Figure 2.**
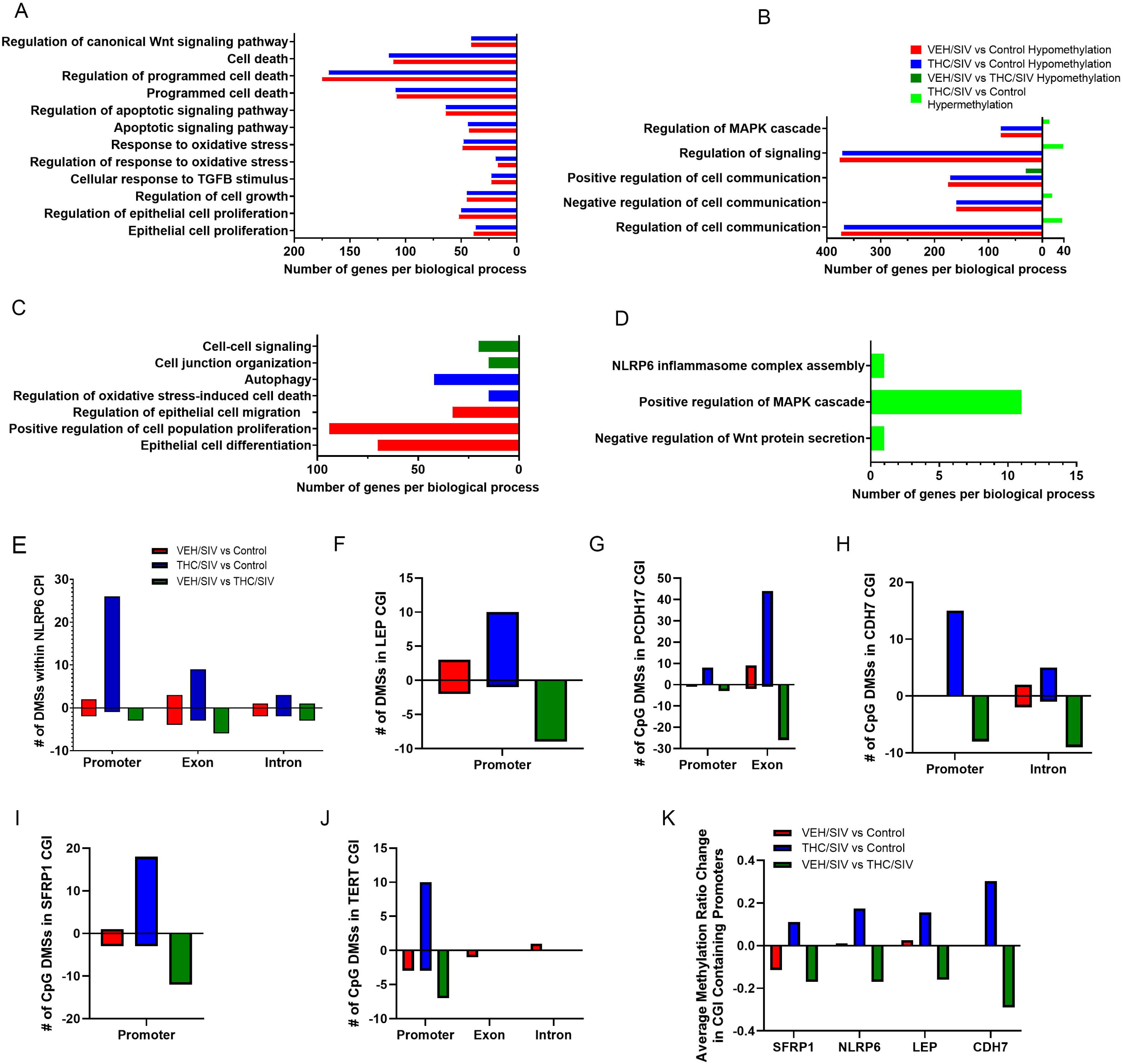
Genes with methylation change throughout CGIs associated with inflammatory response, cellular adhesion, and cell proliferation. Biological processes identified using Gene ontology enrichment analysis of hypomethylated genes in VEH/SIV (red) and THC/SIV (blue) compared to control or in VEH/SIV relative to THC/SIV RMs (gold) (**A**-**C**) or hypermethylated genes in THC/SIV compared to control RMs (green) (**B&D**). Number of differentially methylated CpGs in the CGI of *NLRP6* (**E**), *LEP* (**F**), *PCDH17* (**G**), *CDH7* (**H**), *SFRP1* (**I**), *TERT* (**J**), and the Δβ-values in promoter associated CGIs of *SFRP1*, *NLRP6*, *LEP*, and *CDH7* (**K**). DMSs p-value <0.05, absolute Δβ-value >0.1.

Not surprisingly, compared to controls, only VEH/SIV RMs showed hypomethylation changes in genes involved in epithelial cell differentiation, positive regulation of cell population proliferation, and regulation of epithelial cell migration, an indirect indicator of host response to epithelial cell loss (Figure 2C). Notably, genes responsible for the regulation of oxidative stress-induced cell death and autophagy were hypomethylated in THC/SIV relative to control RMs (Figure 2C). Compared to THC/SIV, VEH/SIV RMs showed hypomethylation of genes involved in cell junction organization and cell-to-cell signaling (Figure 2C). Most strikingly, hypermethylation of genes responsible for the negative regulation of Wnt protein secretion, positive regulation of MAPK cascade, and, more importantly, NLRP6 inflammasome complex assembly (Figure 2D) were detected in THC/SIV relative to control RMs (Figure 3D).

**Figure 3.**
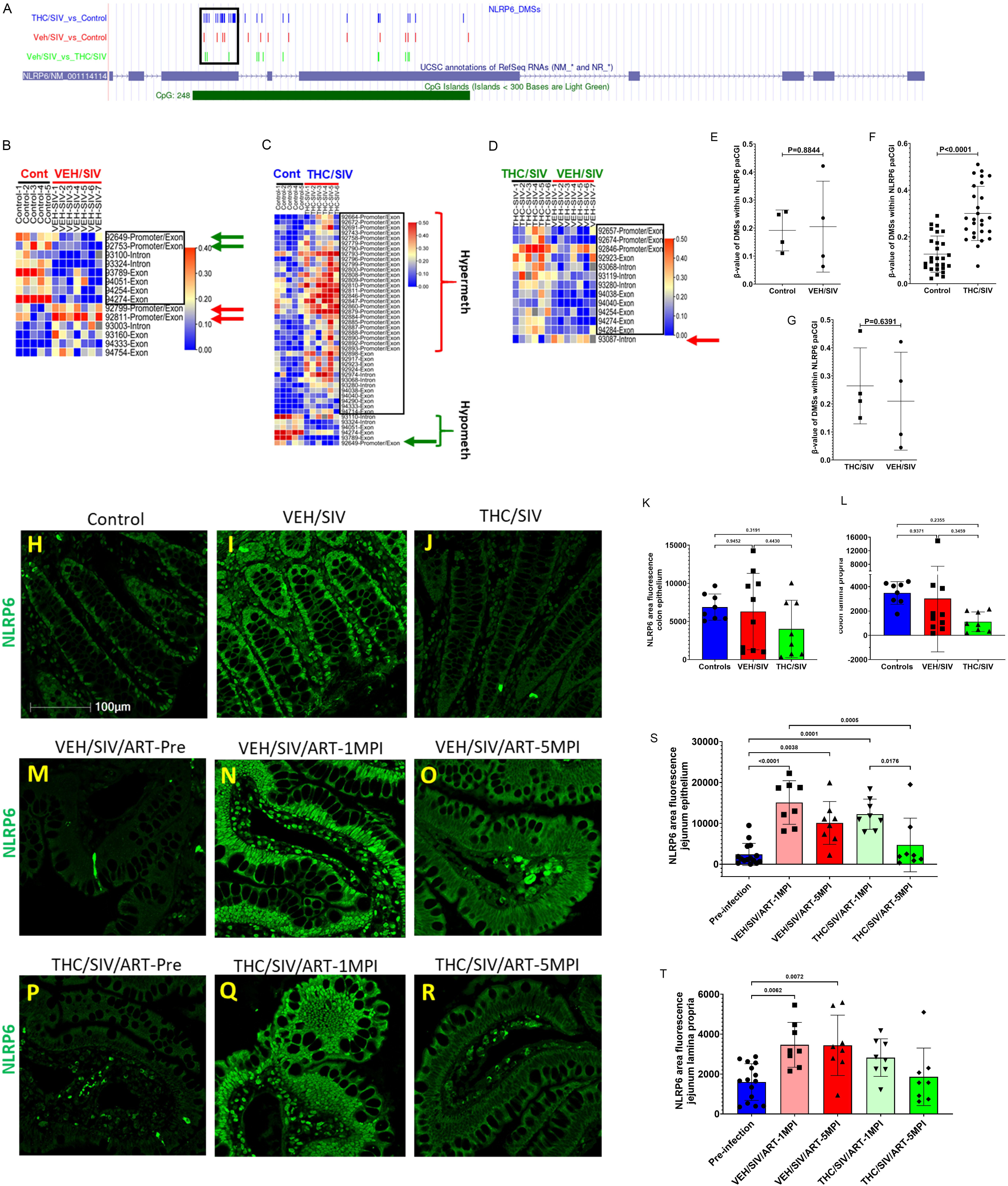
Long-term low-dose THC administration significantly increased methylation in the *NLRP6* promoter associated CGI (paCGI). UCSC Genome track for *NLRP6* showing the location of the paCGI (black box), and DMSs in VEH/SIV (red) and THC/SIV (blue) relative to control, and VEH/SIV compared to THC/SIV RMs (green) (**A**). Heat maps showing the DMSs in NLRP6 CGI in VEH/SIV (**B**), and THC/SIV (**C**) compared to control or in VEH/SIV relative to THC/SIV RMs (**C**). Cumulative β-values of NLRP6 paCGI in VEH/SIV (**E**) and THC/SIV (**F**) relative to control and in VEH/SIV compared to THC/SIV RMs (**G**). Immunofluorescence localization of NLRP6 protein expression in the colon of control (**H**), ART-naïve VEH/SIV (**I**), and THC/SIV RMs (**J**), and jejunum of cART-experienced RMs before SIV infection (**M&P**), at 1 (**N&Q**)- and 5 (**O&R**)-months post infection (MPI) administered Vehicle or THC. NLRP6 protein quantification in colon and jejunum epithelial (**K&S**) and lamina propria mononuclear cell (**L&T**) compartments.

Using a supervised analysis, we examined genes that showed differential methylation throughout their respective paCGIs. Relative to controls, VEH/SIV RMs had DMSs (mostly hypomethylation) in genes associated with inflammatory response [C-C motif chemokine ligand 12 (*CXCL12*), toll-interacting protein (*TOLLIP*), NLR family pyrin domain containing 6 (*NLRP6*; Figure 2E), leptin (*LEP*; Figure 2F), Mab-21 Domain Containing 1 (*MB21D1*) also known as cyclic GMP-AMP synthase (cGAS)], cellular adhesion [claudin 6 and 11, intercellular adhesion molecule 1 (*ICAM1*), protein kinase C-beta (*PRKCB*), protocadherin 17 and 19 (*PCDH17* and *PCDH19*; Figure 2G), adrenocorticotropic receptor beta 1 (*ADRB1*), cadherin 7 (*CDH7*; Figure 2H), genes associated with protection against oxidative stress [cytoglobin (*CYGB*), glutathione peroxidase 3 (*GPX3*)], colonic epithelial cell proliferation [Wnt inhibitory factor 1 (*WIF1*), soluble frizzled protein 1 (*SFRP1*; Figure 2I), telomerase reverse transcriptase (*TERT*; Figure 2J), heart and neural crest derivatives expressed 2 (*HAND2*)], and apoptosis [death-associated protein kinase 1 (*DAPK1*), BCL2-Like-1 (*BCL2L1*)]. Table S1 shows the respective number of DMSs for each gene, as well as the average change in methylation due to these DMSs throughout the CGI. The average methylation changes of 1-14 DMSs in the CGI for the genes above were between −40.7% and 12.3% (columns 4, 6, and 8 of Table S1).

In contrast, relative to control RMs, although THC/SIV RMs showed DMSs in the genes described above, key genes associated with the inflammatory response (Table S1) showed more hypermethylated CpG sites (hyper-MSs), compared to VEH/SIV RMs, specifically *NLRP6* (38 hyper-MSs and 6 hypo-MSs; Figure 2E), and *MB21D1* or *cGAS* (9 hyper-MSs and 1 hypo-MS). The average methylation differences were 12.5% (*NLRP6*) and 24.2% (*MB21D1*). Interestingly, while there were more hyper-MSs in *LEP* (Figure 2F), in THC/SIV than VEH/SIV relative to control RMs, both had an average positive methylation change of 2.6% (VEH/SIV) and 15.6% (THC/SIV). As with *LEP*, while *PCDH17* had more hyper-MSs (52; Figure 2G, Table S1), there was a positive methylation change in both VEH/SIV and THC/SIV compared to control RMs (3.7% and 24.1%, respectively). Genes associated with cellular adhesion (Table S1) showed a similar pattern of methylation difference in THC/SIV and VEH/SIV compared to control RMs, except for *CDH7* (Figure 2H), which had 20 hyper-MSs, one hypo-MS, and a methylation change of 24.9% in THC/SIV compared to control RMs. Like VEH/SIV RMs, genes associated with oxidative stress and apoptosis (Table S1) also showed similar methylation changes in THC/SIV compared to controls. Finally, *SFRP1* (18 hyper-MSs and 3 hypo-MSs; Figure 2I), *TERT* (10 hyper-MSs and one hypo-MS; Figure 2J), and *HAND2* (one hyper-MS), associated with colonocyte proliferation, were hypermethylated in THC/SIV RMs compared to controls, as opposed to VEH/SIV RMs.

Relative to THC/SIV, hypomethylation occurred in all the genes discussed above, with an average methylation difference of between −28 and −11% in VEH/SIV RMs. Since our focus was on methylation changes that occurred in paCGIs, we investigated the average methylation changes that occurred in genes described above that had at least 10% of the CpG sites differentially methylated, which included *NLRP6*, *LEP*, *CDH7*, and *SFRP1* (Figure 2E, 2F, 2H, and 2I, Table S1). Unlike THC/SIV RMs, relative to control RMs, for the genes listed above, VEH/SIV RMs displayed a negative (hypomethylation) methylation change in *TERT and SFRP1* (Figure 2I, 2J and 2K), while *NLRP6* and *LEP* had a positive methylation change (Figure 2K). Interestingly, while no DMSs were detected in the paCGIs of *CDH7* in VEH/SIV compared to control RMs, several DMSs were detected in VEH/SIV relative to THC/SIV RMs.

### Long-term, low-dose THC-induced hypermethylation of CpGs in the paCGI of NLRP6 resulted in decreased NLRP6 protein expression in the colon of SIV-infected RMs

The GO identification of a cluster of hypermethylated genes that were involved in NLRP6 inflammasome assembly (Figure 2D) prompted us to focus further on methylation changes in *NLRP6*, the gene responsible for protecting the intestinal epithelial barrier from harmful gut microbes and maintaining gut homeostasis, through inflammasome activation. However, in IBD, although the host responds to microbial invasion initially by activating NLRP6, the persistence of the inflammatory trigger drives NLRP6 hyperactivation, inflammasome activation, and consequently damaging colonic inflammation and barrier disruption ^45^. Within the *NLRP6* CGI, VEH/SIV RMs showed a total of 14 DMSs (Figures 3A and 3B), relative to controls. These included eight hypomethylated (boxed region in Figure 3B) and six hypermethylated sites. Within the promoter region, two sites each were hypomethylated (green arrows in Figure 3B) and hypermethylated (red arrows in Figure 3B). Relative to controls, THC/SIV RMs showed a significantly higher number of DMSs in *NLRP6* CGI than seen in VEH/SIV RMs, with a total of 44 DMSs (Figure 3A and 3C). Out of these, 38 sites were hypermethylated (boxed region in Figure 3C), and 6 were hypomethylated. Strikingly, except for one (hypomethylated) (green arrow in Figure 3C), 26 sites were hypermethylated (red curly bracket in Figure 3C) in the paCGI region. Lastly, when compared to THC/SIV, VEH/SIV RMs had 13 DMSs in the CGI of NLRP6 (Figure 3A and 3D), out of which 12 were significantly hypomethylated (boxed region in Figure 3D) and only one site located in the intron was hypermethylated (red arrow in Figure 3D). The average Δβ-value in the paCGI of NLRP6 was not significantly different in VEH/SIV relative to controls (Figure 3E) or THC/SIV RMs (Figure 3G). Inversely, there was a significant increase in the Δβ-value of CpG sites of *NLRP6* paCGI in THC/SIV compared to control RMs (Figure 3F).

Since methylation changes alter gene expression, we next investigated NLRP6 protein expression levels in the colon of all three groups. Since microarray data did not detect significant changes in *NRLP6* mRNA expression, we focused on NLRP6 protein expression because not all genes show similar changes at the mRNA and protein levels. NLRP6 protein expression was detected in both the nucleus and cytoplasm, with a stronger signal in the nucleus of CE and lamina propria mononuclear cells (LPMNCs). We found an inverse relationship between hypermethylation changes in the paCGI of NLPR6 and protein expression, although NLRP6 staining intensity was not significantly different in VEH/SIV RMs (n=6; Figure 3I and 3K) or THC/SIV RMs (n=6; Figure 3J and 3K) compared to controls (n=5; Figure 3H and 3K). Nevertheless, NLRP6 staining intensity had a decreasing trend exclusively in CE of THC/SIV RMs (Figure 3J and 3K) relative to controls and VEH/SIV RMs. Similar to the CE, NLRP6 protein expression was reduced in LPMNCs (Figure 3L).

### Acute HIV/SIV infection significantly increased NLRP6 protein expression, while low-dose THC in combination with cART successfully decreased its expression in the jejunum of SIV-infected RMs

Next, we looked at the effect of acute HIV/SIV infection [1 month post infection (MPI)] on NLRP6 protein expression and if cART alone or in combination with THC could decrease NLRP6 protein expression. Here, we focused on the jejunum, as it plays a critical role in both nutrient and cART drug absorption, and its dysfunction can negatively impact both functions. Moreover, high NLRP6 protein expression has been confirmed in the jejunum of mice ^46^ and humans ^47,48^. Longitudinally, acute HIV/SIV infection (1MPI; n=8/group; Figure 3N, 3Q, and 3S) resulted in a significant increase in NLRP6 protein expression compared to pre-infection (Figure 3M, 3P, and 3S). At 5MPI, NLRP6 expression in cART-suppressed (VEH/SIV/cART; n=8) RMs did not differ from their 1MPI timepoint but significantly elevated from the pre-infection timepoint (Figure 3O, 3N, 3M, and 3S). Strikingly, THC/SIV/cART RMs (n=8) showed a significant decrease in NLRP6 protein expression from their respective 1MPI timepoints (Figure 3R, 3Q, and 3S) but showed no significant decrease from VEH/SIV/cART RMs at 5MPI or the pre-infection timepoint (Figure 3O, 3P, and 3S). THC-induced reductions in NLRP6 protein expression in the jejunum LPMNCs showed a similar trend (Figure 3T) to that observed in the epithelium (Figure 3S).

### Phytocannabinoids blocked polyI:C and lipoteichoic acid-induced NLRP6 protein upregulation in vitro and significantly increased mRNA expression of the NLRP6 deubiquitinating enzyme CYLD in the CE in vivo

To determine if phytocannabinoids directly blocked NLRP6 protein expression, we separately added 3 µM THC and CBD to human small intestinal epithelial cultures after transfection with 2µg/mL of polyI:C followed by treatment with lipoteichoic acid (LTA) (10µg/mL) 4 h later. It is clear from Figure 4B, 4C, and 4I that polyI:C alone or in combination with LTA significantly increased NLRP6 protein expression in the cytoplasm (white arrows in Figure 4B and 4C) compared to cells treated with DMSO (Figure 4A and 4I). Interestingly, pretreatment for 1 h with THC (Figure 4D and 4I) or CBD (Figure 4G and 4I) or both (Figure 4H and 4I) successfully blocked polyI:C and LTA-induced NLRP6 protein upregulation. Treatment of cells with an antagonist of the cannabinoid receptor-1 (AM251) (Figure 4E and 4I) or cannabinoid receptor-2 (AM630) (Figure 4F and 4I) did not alter THC’s ability to counter polyI:C and LTA-induced NLRP6 protein upregulation. Therefore, future studies are needed to determine the role of other receptors such as GPR55, PPARs and TRPV ion channels in cannabinoid blockade of NLRP6 protein induction.

**Figure 4.**
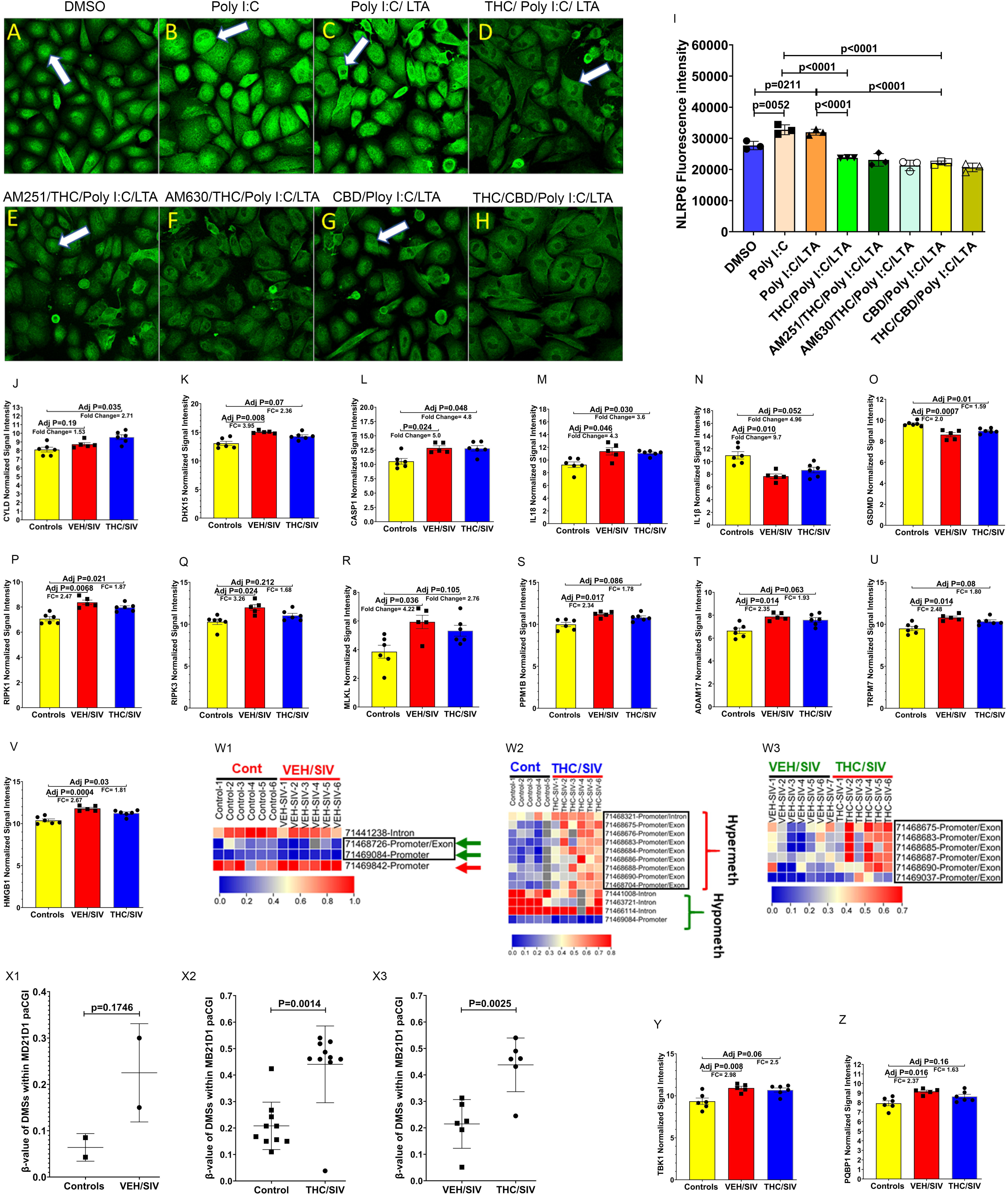
Cannabinoids successfully blocked polyI:C and LTA-induced NLRP6 upregulation in vitro and reduced necroptosis-driven gene signature in colonic epithelium (CE) of SIV-infected rhesus macaques (RMs). Representative immunofluorescence images showing the expression of NLRP6 (green) protein in human small intestinal epithelial (HSIE) cells at 18 h following treatment with DMSO (**A**) or post transfection with 2 µg/mL of polyI:C alone (**B**) or also treated with lipoteichoic acid (LTA) (10 µg/mL) 4 h later (**C**). PolyI:C and LTA treated HSIE cells were preincubated for 1 h with THC (**D**), AM251 + THC (**E**), AM630 + THC (**F**), CBD (**G**) or THC and CBD **(H**), fixed after 18 h, stained and NLRP6 (green) protein expression quantitated (**I**). Experiments were performed in triplicate wells using 3 µM of THC and CBD, 10 µM of AM251/AM630, and repeated thrice. Microarray normalized signal intensity of *CYLD* (**J**), *DHX15* (**K**), *CASP1* (**L**), *IL-18* (**M**), *IL-1*β (**N**), *GSDMD* (**O**), *RIPK1* (**P**), *RIPK3* (**Q**), *MLKL* (**R**), *PPM1B* (**S**), *ADAM17* (**T**), *TRPM7* (**U**), *HMGB1* (**V**), *TBK1* (**Y**) and *PQBP1* (**Z**) in CE of SIV-infected RMs. Heat maps showing the DMSs in MB21D1 CGI in VEH/SIV (**W1**), and THC/SIV (**W2**) compared to control or in VEH/SIV relative to THC/SIV RMs (**W3**). Cumulative β-values of MB21D1 paCGI in VEH/SIV (**X1**) and THC/SIV (**X2**) relative to control and in VEH/SIV compared to THC/SIV RMs (**X3**). White arrows in panels A to G indicate epithelial regions. Immunofluorescence data were analyzed using one-way ANOVA followed by Tukey’s multiple comparison post hoc test. A p-value of <0.05 was considered significant.

Next, we examined if the gene expression levels of the *NLRP6* inflammasome components and proinflammatory cytokines changed consequent to reduction of *NLRP6* protein expression in response to hypermethylation of the paCGI. Intriguingly, mining our recently published CE microarray data from the same RMs ^44^ for genes that showed differential expression by at least 1.5-fold change and an adjusted p≤0.05 confirmed statistically significant upregulation of *CYLD*, a deubiquitinase shown to remove K63-linked ubiquitin residues on NLRP6 ^45^ and attenuate its activity. *CYLD* mRNA was significantly upregulated in CE of THC/SIV (∼2.7-fold) but not VEH/SIV RMs relative to controls (Figure 4J). Similarly, *DHX15*, an RNA helicase and double-stranded viral RNA sensor shown to interact with NLRP6 and trigger NLRP6 inflammasome assembly and activation ^49^, was significantly upregulated in CE of VEH/SIV but not in THC/SIV RMs relative to controls (Figure 4K). In addition, while mRNA expression of caspase-1 (*CASP1*, Figure 4L) and interleukin-18 (*IL-18*, Figure 4M) increased significantly, the opposite trend (downregulated) was detected with interleukin-1β (*IL-1*β, Figure 4N) in CE of SIV-infected RMs irrespective of VEH or THC treatment relative to controls. Finally, mRNA expression of the membrane pore-forming protein, Gasdermin D (*GSDMD*), the downstream executor of pyroptosis, was significantly reduced in CE of both VEH/SIV and THC/SIV relative to control RMs (Figure 4O).

### THC-induced NLRP6 promoter hypermethylation associated with reduced necroptosis-driving gene signature in the CE of SIV-infected RMs

Although *CASP1* and *IL18* mRNA expression was significantly upregulated in CE of both VEH/SIV and THC/SIV RMs, the significantly decreased *GSDMD* mRNA expression in both groups suggested that pyroptosis may not be the predominant pathway driving CE dysfunction. Nevertheless, *NLRP6* has been shown to increase the expression of mixed lineage kinase domain like pseudokinase protein (*MLKL*) and receptor-interacting serine/threonine-protein kinase-3 (*RIPK3*) to induce necroptosis ^50^, a caspase-independent inflammatory mode of cell death. Interestingly, again mining our recently published microarray data ^44^ for genes that showed differential expression by at least 1.5-fold change and an adjusted p≤0.05 confirmed statistically significant upregulation of *RIPK1*, *RIPK3*, and *MLKL* in CE of VEH/SIV RMs (Figure 4P-R). While expression of *RIPK1* mRNA showed statistically significant increase in both groups (Figure 4P), mRNA expression of *RIPK3* and *MLKL* showed statistically significant increase only in CE of VEH/SIV (Figure 4Q and 4R) compared to control RMs. Moreover, *RIPK3*, phosphorylated on Ser227 and Ser199, plays a crucial role in the activation of *MLKL* by phosphorylating it on Thr357 and Ser358 to enable its loading onto the necrosome ^51^. In this context, *PPM1B*, a phosphatase known to dephosphorylate *RIPK3* ^51^ was also significantly upregulated only in CE of VEH/SIV RMs (Figure 4S), indirectly suggesting the presence of p-RIPK3. Most strikingly, mRNA expression of *ADAM17* (Figure 5T), a positive regulator of necroptosis ^52^, *TRPM7* (Figure 5U), a downstream target of *MLKL* that promotes its membrane localization and Ca^2+^ ion influx for initiating necroptosis ^53^, and *HMGB1* (Figure 5V), a danger-associated molecular pattern (DAMP) protein, known to propagate inflammation associated with both pyroptotic and necroptotic cell death ^54^ was significantly increased only in CE of VEH/SIV RMs.

**Figure 5.**
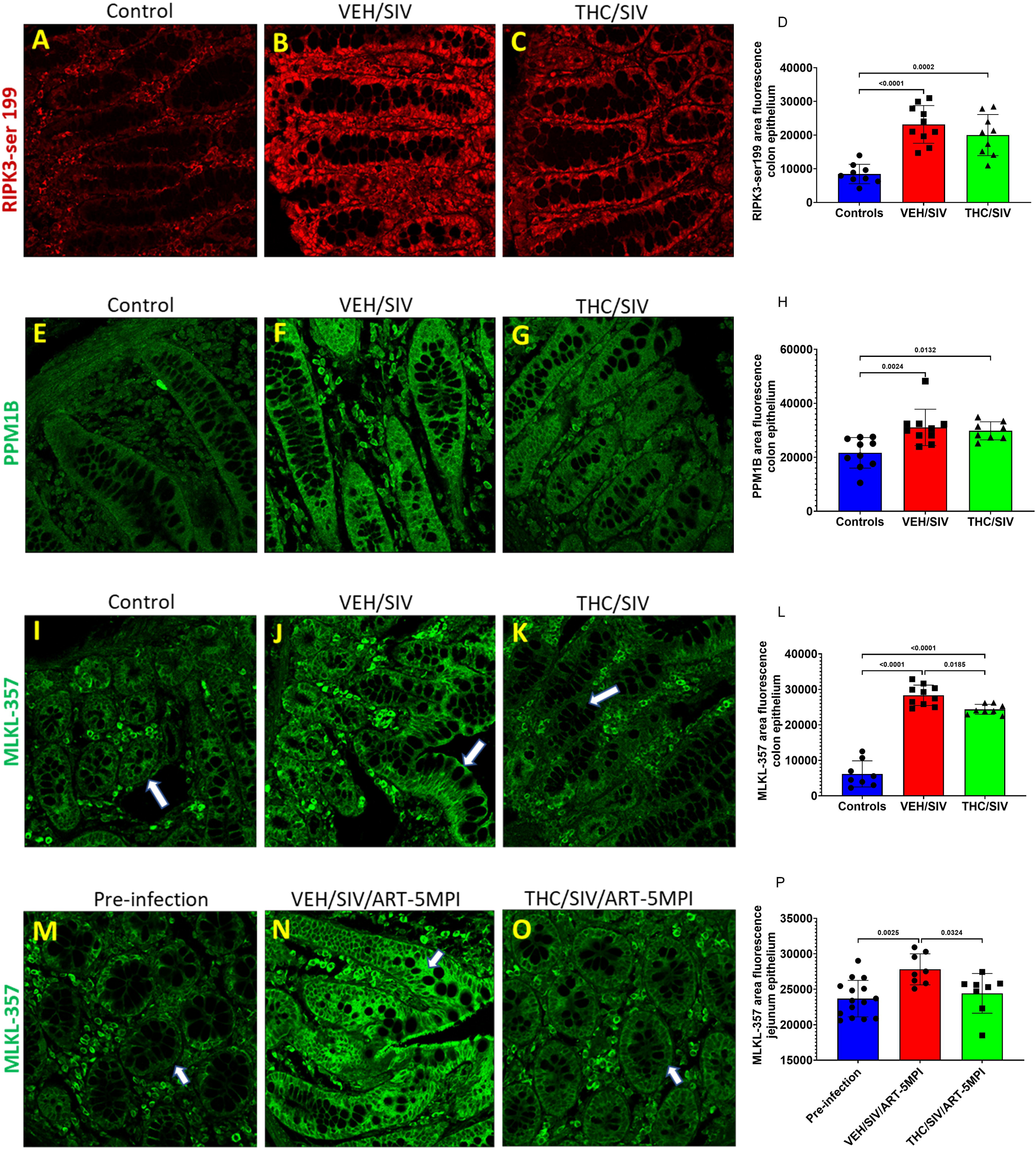
Chronic THC administration reduced HIV/SIV infection induced RIPK3(Ser199), PPM1B, p-MLKL(Thr357) protein expression in the intestine. p-RIPK3(Ser199) (red) (**A-C**), PPM1B (**E-G**) (green) and p-MLKL(Thr357) (green) (**I-K**) protein expression in colon tissues of uninfected control (**A, E&I**), VEH/SIV (**B, F&J**), and THC/SIV rhesus macaques (RMs) (**C, G&K**). p-MLKL(Thr357) (green) protein expression in jejunum tissues before SIV infection (**M**) and at 5 months post SIV infection in cART suppressed VEH/SIV (**N**) and THC/SIV (**O**) RMs. Representative immunofluorescence images were captured using a Zeiss confocal microscope at 20X magnification. Quantitation of p-RIPK3(Ser199) (**D**), PPM1B (**H**) and p-MLKL(Thr357) (**L&P**) average positive area fluorescence in colon and jejunum epithelium was performed using Halo software. Note the significantly reduced p-MLKL(Thr357) staining in the colon and jejunum epithelium of THC/SIV (**K, O, L&P**) compared to VEH/SIV (**J, N, L &P**) RMs. White arrows in panels **I** to **O** indicate epithelial regions. Immunofluorescence data were analyzed using one-way ANOVA followed by Tukey’s multiple comparison post hoc test. A p-value of <0.05 was considered significant.

We previously reported significantly reduced interferon (IFN)-stimulated gene expression (ISG) in whole colon ^39^, CE ^44^ and basal ganglia (brain) ^8^ of THC/SIV RMs. Nevertheless, the mechanism of THC action remained unknown. *MB21D1,* or *cGAS,* is an important DNA sensor that drives type-I IFN responses ^55^. In this context, within the *MB21D1* CGI (Figure S3A), VEH/SIV RMs showed a total of 4 DMSs (Figure 4W1), relative to controls. These included three hypomethylated (Figure 4W1) and one hypermethylated site. Within the paCGI, two sites were hypomethylated (green arrows in Figure 4W1) and one hypermethylated (red arrow in Figure 5W1). Relative to controls, THC/SIV RMs showed a significantly higher number of DMSs in *MB21D1* CGI than seen in VEH/SIV RMs, with a total of 13 DMSs (Figure 4W2). Out of these, nine sites, all in the paCGI region, were hypermethylated (boxed region and red curly bracket in Figure 4W2), and four were hypomethylated (green curly bracket in Figure 4W2). Lastly, when compared to THC/SIV, VEH/SIV RMs had six DMSs in the CGI of *MB21D1* (Figure 4W3), all of which were in the paCGI region (boxed region in Figure 4W3). The average Δβ-value in the paCGI of *MB21D1* was not significantly different in VEH/SIV relative to controls (Figure 4X1). In contrast, there was a significant increase in the Δβ-value of CpG sites of *MB21D1* paCGI in THC/SIV compared to control (Figure 4X2) and VEH/SIV (Figure 4X3) RMs. Further, the expression of *TBK1*, a signaling adapter protein that transduces the signal emanating from *MB21D1* ^56^, was significantly increased in CE of VEH/SIV but not THC/SIV RMs (Figure 4Y). Finally, *PQB1*, a proximal sensor of *MB21D1*-dependent increase in type-I IFN response to HIV-1 infection ^57^, was also significantly increased in CE of VEH/SIV but not THC/SIV RMs (Figure 4Z).

### Protein expression of necroptosis mediators p-RIPK3 (Ser-199), p-MLKL (Thr-357/Ser-358), and HMGB1 is significantly increased in the intestinal epithelium of SIV-infected RMs

Activated RIPK1 promotes RIPK3 phosphorylation, which then autophosphorylates itself on Ser199 (required for kinase function) and Ser227 (required for necroptosis), two sites crucial for the activation of MLKL. Interestingly, p-RIPK3(Ser199) showed significantly higher expression in CE of both ART-naïve VEH/SIV and THC/SIV relative to control RMs (Figure 5A to 5D) but not in ART-experienced RMs (Figure S3B-3E). In this context, protein expression of PPM1B, a phosphatase that inactivates RIPK3 by dephosphorylation, was significantly increased in CE of both ART-naïve VEH/SIV and THC/SIV relative to control RMs (Figure 5E to 5H) but not in ART-experienced RMs (Figure S3F-3I). The high expression of PPMB1 indirectly indicates the presence of activated p-RIPK3. Note that for both p-RIPK3(Ser199) and PPM1B, protein expression was slightly higher in VEH/SIV than THC/SIV relative to control RMs.

Consistent with high *MLKL* mRNA expression in CE of VEH/SIV RMs, protein expression of p-MLKL(Thr357), a marker of necroptosis, was significantly upregulated with intense cytoplasmic expression in CE of ART-naïve (Figures 5J and 5L) and ART-experienced VEH/SIV (Figure 5N and 5P) relative to control (Figures 5I and 5M) RMs. p-MLKL(Thr357) expression in CE of THC/SIV RMs was significantly higher than controls (Figures 5K and 5L). Most notably, p-MLKL(Thr357) protein expression was significantly reduced in CE of THC/SIV compared to VEH/SIV RMs in both ART-naïve (Figure 5K and 5L) and ART-experienced RMs (Figure 5O and 5P).

Likewise, p-MLKL(Ser358) protein expression was significantly increased in CE of both VEH/SIV and THC/SIV relative to control RMs (Figures 6A to 6D) in CE. Similar to p-MLKL(Thr357), protein expression of p-MLKL(Ser358) was more intense in the cytoplasm of VEH/SIV but not in THC/SIV compared to control RMs (Figure 6B and 6C). Notably, at 5MPI, p-MLKL(Ser358) protein expression was significantly reduced in THC/SIV/ART compared to VEH/SIV/ART RMs (Figure 6F, 6G, and 6H). While no significant differences in protein expression of p-RIPK3(Ser199) were detected in the jejunum of SIV/ART RMs, it is possible that p-RIPK3(Ser227) ^51^ and the Tam family of receptor tyrosine kinases ^58^ may be the predominant kinases phosphorylating MLKL in the jejunum. Finally, expression of HMGB1, an important DAMP released upon pyroptotic and necroptotic cell death and an initiator of secondary inflammatory responses in neighboring cells^59^, showed a marked but statistically nonsignificant reduction in CE of ART-naïve THC/SIV RMs compared to VEH/SIV RMs (Figure 6I to 6L). Nevertheless, unlike colon, protein expression of HMGB1 was significantly increased in the jejunum epithelium of VEH/SIV/ART RMs compared to pre-infection and THC/SIV/ART RMs (Figure 6M to 6P). Note the restoration of HMGB1 protein expression to pre-infection levels by low-dose THC at 5MPI (Figure 6P). These findings demonstrate, for the first time, the activation of necroptosis and its contribution to intestinal dysfunction in HIV/SIV infection under suppressive cART and provide novel clinically relevant insights into how low-dose phytocannabinoids, potentially through epigenetic modulation of NLRP6, can reduce necroptosis signaling potentially via modulation of the RIPK3-MLKL pathway (Figure 7) that may help reduce intestinal inflammation in PLWH and, by extension, IBD patients.

**Figure 6.**
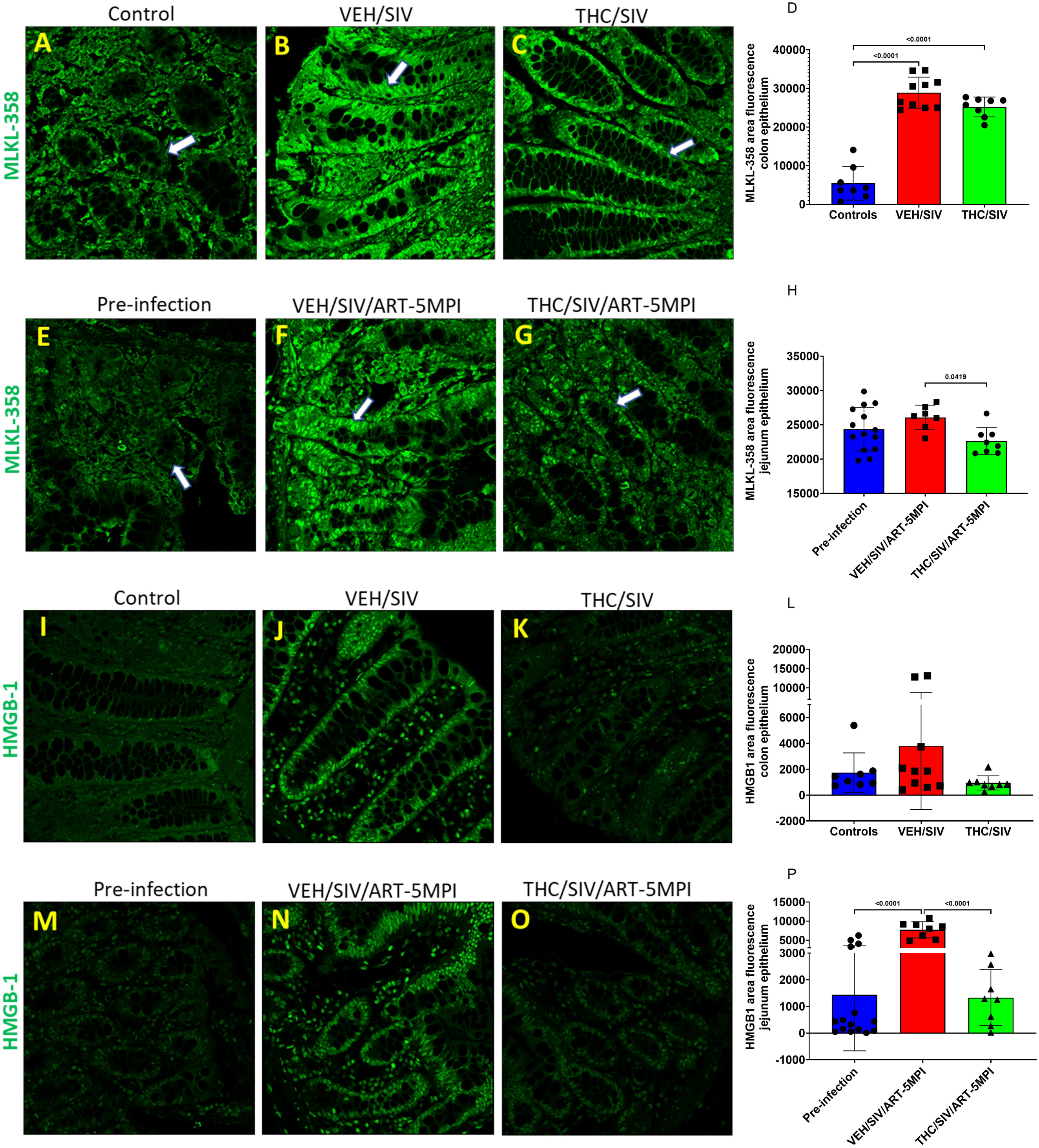
Chronic THC administered in combination with cART significantly reduced HIV/SIV infection induced p-MLKL(Ser358) and HMGB1 protein expression in the intestine. p-MLKL(Ser358) (green) (**A-C**), and HMGB1 (green) (**I-K**) protein expression in colon tissues of uninfected control (**A&I**), VEH/SIV (**B&J**), and THC/SIV (**C&K**) rhesus macaques (RMs). p-MLKL(Ser358) (green) (**E-G**) and HMGB1 (green) (**M-O**) expression in jejunum tissues before SIV infection (**E&M)**) and at 5 months post SIV infection in cART suppressed VEH/SIV (**F&N**) and THC/SIV (**G&O**) RMs. Representative immunofluorescence images were captured using a Zeiss confocal microscope at 20X magnification. Quantitation of MLKL(Ser358) (**D&H**), and HMGB1 (**L&P**) average positive area fluorescence in colon and jejunum epithelium was performed using Halo software. Note the significantly reduced p-MLKL(Ser358) and HMGB1 staining in the jejunum epithelium of cART suppressed THC/SIV (**G, O, H&P**) compared to VEH/SIV (**F, N, H&P**) RMs. White arrows in panels A to G indicate epithelial regions. Immunofluorescence data were analyzed using one-way ANOVA followed by Tukey’s multiple comparison post hoc test. A p-value of <0.05 was considered significant.

**Figure 7.**
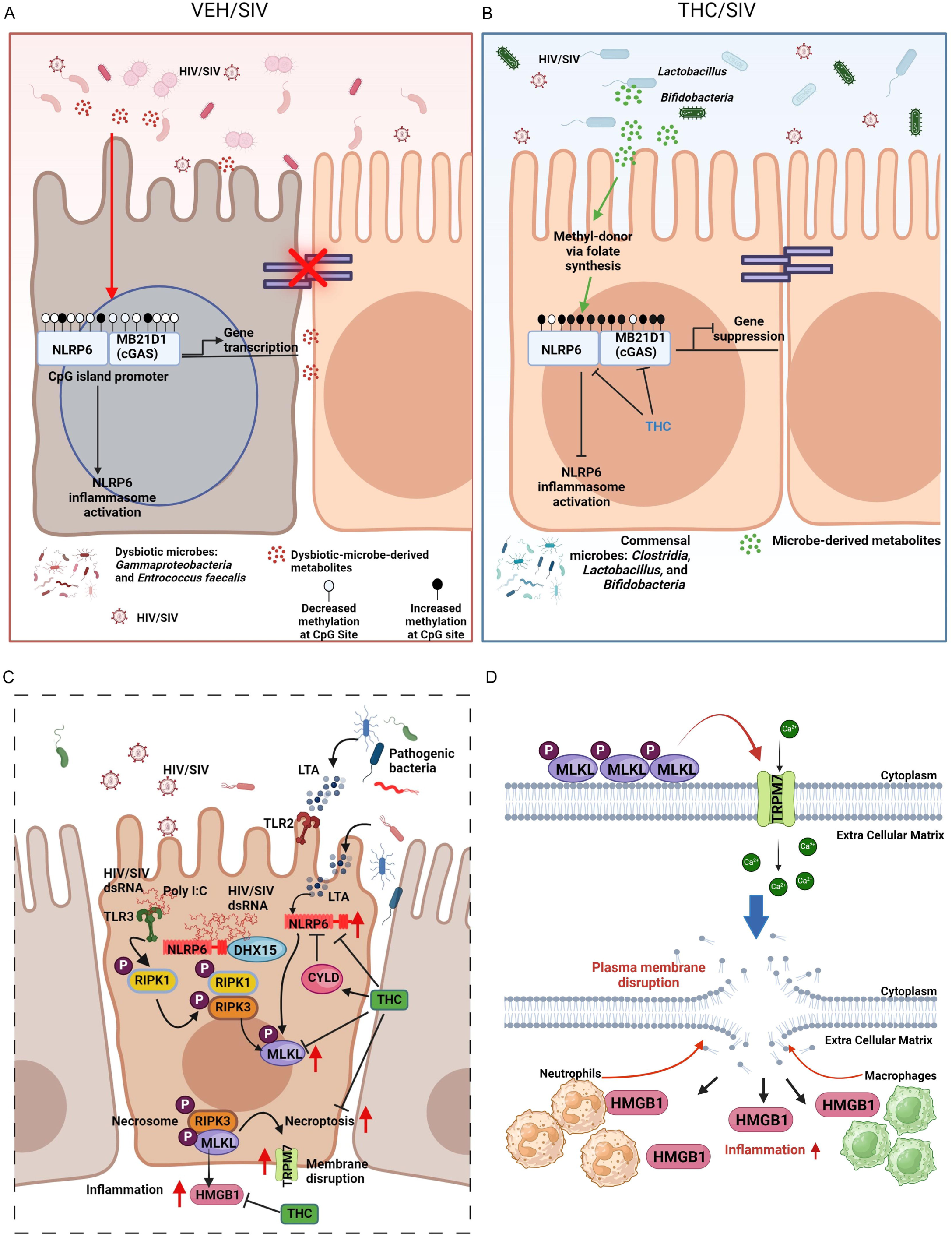
HIV/SIV infection-induced DNA hypomethylation facilitates persistent activation of the NLRP6 sensor and cGAS/MB21D1 protein expression in the intestinal epithelium, leading to epithelial barrier breakdown and microbial translocation. (**A**). Long-term low-dose THC epigenetically modulates these changes by hypermethylating CpG sites in the NLRP6 and cGAS promoters, thereby reducing gastrointestinal mucosal injury (**B**). NLRP6 inflammasome assembly could potentially activate the RIPK1-RIPK3-MLKL pathway to induce necroptosis of intestinal epithelium (**C**). TLR3 (Figure S3J) and NLRP6-DHX15 viral sensing complexes could recognize dsRNA intermediates of HIV, leading to the phosphorylation of RIPK1, which in turn activates RIPK3, culminating in the activation of the necroptosis executioner protein MLKL through phosphorylation on Thr357 and Ser358. Activation of MLKL and concurrent activation of the calcium ion channel TRPM7 enhances intracellular Ca++ influx, leading to necroptotic cell death. Cell death facilitates the release and extracellular accumulation of the DAMP protein, HMGB1, which can propagate secondary inflammatory responses by attracting neutrophils and macrophages leading to more widespread epithelial damage (**D**). By hypermethylating the NLRP6 promoter and increasing expression of the deubiquitinase CYLD, long-term low-dose THC can block the downstream activation of the RIPK1-RIPK3-MLKL pathway and significantly reduce HMGB1 protein expression to attenuate gastrointestinal inflammation and preserve epithelial barrier integrity. Figures were generated using Biorender.

## Discussion

Intestinal epithelial barrier disruption, triggered by mucosal immune dysfunction, is a common sequela to HIV/SIV infection despite peripheral viral suppression by cART, which leads to systemic inflammation, a major contributor to HIV-associated comorbidities ^1^. Persistent intestinal inflammation can trigger and sustain the accumulation of epigenetic modifications, particularly alterations in levels of promoter DNA methylation in genes linked to epithelial proliferation ^27^ leading to enterocyte senescence, epithelial barrier dysfunction, and dysbiosis ^60^. Nevertheless, epigenetic control of epithelial barrier function in HIV/SIV infection remains an understudied topic. In this context, chronic intestinal inflammation and epigenetic alterations involving DNA methylation in gene promoters, together with transcriptome changes in intestinal epithelial cells, have been associated with IBD pathogenesis ^61^. Moreover, since epigenetic signatures are cell type-specific, separating out enough purified epithelial cells from human colorectal pinch biopsies (maximum n=3 biopsies) is challenging. The feasibility of obtaining intact colon resections (∼5 cm) without any adverse effects from SIV-infected RMs allows the isolation of ∼85-95% pure epithelial cells and offers unique avenues to identify epigenetically altered genes involved in critical GI mucosal abnormalities. While we found significant global hypomethylation throughout the whole genome during chronic HIV/SIV infection, chronic low-dose phytocannabinoid (THC) administration caused significantly more hypermethylation and effectively increased methylation in paCGIs of genes associated with the inflammatory response (*NLRP6, MB21D1, LEP*), cellular adhesion (*CDH7*), and colonic epithelial cell proliferation (*SFRP1* and *TERT*).

HIV/SIV infection resulted in a significant decrease in methylation levels throughout the genome and specifically in paCGIs. Strikingly, chronic low-dose THC administration caused significantly more changes in methylation, both hypo- and hyper- methylation, than that observed in VEH/SIV RMs compared to controls. Remarkably, hypomethylation of just one DMS in the paCGI had an influence on gene expression (upregulation). This finding leads us to speculate that the traditional idea of methylation being inversely correlated to gene expression is gene location specific. While the paCGIs are regulatory regions, methylation changes in enhancer regions might have more influence on gene expression than paCGIs ^62,63^. Interestingly, methylation changes that occurred outside of a gene region or mapped to an unannotated region were found to be in gene enhancers, which potentially could influence gene expression ^63^. Since our analysis only looked for DMSs within a CGI, and if it was in a gene promoter, exon, or intron, we could not determine if the ∼86% of unannotated sites excluded from the analysis were a part of gene enhancers and the genes they regulated. Differential methylation of enhancers has been implicated in osteoarthritis, Alzheimer’s disease (AD), and multiple sclerosis ^64^. Therefore, further work to identify the unannotated DMSs that are an integral part of gene enhancers with potential role(s) in HIV/SIV-induced intestinal dysfunction is needed.

Significant hypomethylation detected in the paCGIs of genes associated with inflammation, cellular adhesion, oxidative stress, cell proliferation, and apoptosis in CE of VEH/SIV RMs suggests that alterations in the epigenetic landscape may facilitate their increased expression, leading to the persistence of a proinflammatory environment. These findings are consistent with increasing evidence pointing towards an association between DNA hypomethylation and increased expression of proinflammatory genes (76-78), leading to oxidative DNA damage, epithelial cellular senescence, barrier dysfunction, microbial translocation, and systemic inflammation. Interestingly, compared to controls, THC/SIV RMs had significantly more hypermethylation in the paCGI region of genes associated with inflammatory responses, cellular adhesion, and colonic epithelial cell proliferation, specifically *NLRP6*, *CDH7*, *SFRP1*, and *TERT*. These genes showed increased methylation levels in CE of THC/SIV compared to control RMs that accounted for ∼4-23% of CpG sites in their respective CGIs, and the average methylation difference from all methylation changes in these genes was ∼18%. More importantly, THC hypermethylated promotor CpG sites of *MB21D1*, also known as *cGAS*, which interacts with *PQBP1* following its binding to retroviral reverse transcribed DNA in the cytosol to activate the cGAS-STING pathway, leading to increased type-I interferon (IFN) production. Interestingly, in addition to promoter CpG hypermethylation of *MB21D1*, *PQBP1* gene expression was also significantly reduced in the CE of THE/SIV RMs, which may partly explain the reduced type-I interferon gene expression in the CE of THC/SIV RMs we reported previously ^39,44^, suggesting that phytocannabinoids are effective in inhibiting cGAS-STING signaling, a pathway hyperactivated in ulcerative colitis leading to dysregulated intestinal epithelial cell integrity, autophagy, and immune/inflammatory responses ^55^. Interestingly, gene ontology analysis identified significant hypomethylation of genes that regulate oxidative stress-induced cell death and autophagy in CE of THC/SIV RMs. While cannabinoids are well-known to induce autophagy pathways to reduce chronic inflammatory responses ^65^, our novel findings demonstrate the involvement of epigenetic mechanisms underlying the enhancement of autophagy (promoter DNA hypomethylation) via reduction of *MB21D1*-mediated type-I IFN expression (promoter DNA hypermethylation) by cannabinoids. Thus, epigenetic modulation represents a key mechanism by which phytocannabinoids alter inflammatory signaling/response, epithelial cell adhesion and proliferation, and autophagy in HIV/SIV infection.

When focusing on epigenetic changes in genes that regulate epithelial barrier maintenance and homeostasis, inflammasome components and sensors, *NLRP6*, stood out prominently. A critical function of *NLRP6* is to protect the intestinal epithelial barrier from invading microbes or viruses, where it shows the highest expression ^66,67^. *NLRP6* is thought to regulate intestinal microbiome homeostasis through its ability to activate other proinflammatory and antimicrobial genes ^46,50,66,68^. In return, the gut microbiome modulates NLRP6 inflammasome through their microbial metabolites, prompting *NLRP6* to activate the production of antimicrobial peptides by intestinal epithelial cells (paneth cells) or triggering goblet cells to induce mucin exocytosis ^66, 48^. While *NLRP6* has been shown to maintain intestinal homeostasis, its perpetual activation due to inadequate regulation, as previously shown in active ileal Crohn’s disease (CD) (131-fold high) and colonic CD patients (3.9-fold high) ^69^, can exacerbate inflammation, leading to gastrointestinal disease and even cancer. Quite unexpectedly, we did not see a change in *NLRP6* mRNA expression in either SIV-infected RM group compared to controls or to each other. Aberrant hypomethylation of *NLRP6* can expedite inflammasome assembly, thereby activating inflammatory signaling pathways leading to CE apoptosis, as seen in Kawasaki disease (KD). Increased *NLRP4*, *NLRP12*, and *IL-1β* gene expression in KD patients was accompanied by hypomethylation of *NLRP4* and *NLRP12* ^70^. Interestingly, treatment with intravenous immunoglobulin restored the gene expression and methylation levels of *NLRP4*, *NLRP12*, and *IL-1β*. In the present study, we show that chronic SIV-infection in cART-naïve RMs neither caused significant hypomethylation changes to *NLRP6* paCGI (2 CpG sites compared to controls and 3 CpG sites compared to THC/SIV RMs) nor increased protein expression significantly in CEs, compared to controls. Nevertheless, hypomethylation of six other CpG sites in the exons and introns may impact *NLRP6* expression. Contrariwise, chronic administration of low-dose THC to chronically SIV-infected RMs resulted in a significant hypermethylation of the *NLRP6* paCGI compared to controls, which resulted in markedly decreased *NLRP6* protein expression in the epithelium and lamina propria cells compared to both control and VEH/SIV RMs.

The decrease in NLRP6 protein expression through hypermethylation was an interesting finding that led us to question the effect of cART on *NLRP6* expression. During acute HIV/SIV infection (1MPI), there was a significant increase in NLRP6 protein expression in the jejunum epithelium, with cART initiated 2 weeks post infection. Noteworthy, at the chronic stage of HIV/SIV infection (5MPI), after viral suppression for ∼4.5 months, cART alone had no effect on NLRP6 expression. However, and most importantly, phytocannabinoid administration, in combination with cART, resulted in significantly decreased NLRP6 protein expression in chronically infected RMs, bringing NLRP6 protein expression down to pre-infection levels. The significantly high NLRP6 protein expression detected in the jejunum of THC/SIV/cART RMs at 1MPI despite initiating THC before SIV infection suggests that phytocannabinoids may produce better anti-inflammatory effects when administered in combination with cART. The successful lowering of NLRP6 protein expression at 5MPI also demonstrates that cART alone, regardless of undetectable viral loads in plasma and tissues, is not sufficient to restore intestinal homeostasis, but long-term, low-dose phytocannabinoids represent an excellent option to be used as an adjunct to cART to reduce damaging intestinal inflammatory responses that persist in PLWH. Folate produced by *Lactobacillus* and *Bifidobacteria*, also shown to be reduced in IBD can affect host cellular DNA methylation, by regulating the methyl donor availability. The increased methylation levels detected in THC/SIV RMs may be driven by the increased relative abundance of *Lactobacillus* (n=28) and *Bifidobacterium* (n=18) species we recently demonstrated ^8^, both of which are commensal microbes that synthesize folate for methyl group donation ^71^. As emphasized by Xu *et al*. ^61^ and Allen *et al.* ^72^, the interaction between gut microbes and host epigenetics should not only be studied further but also therapeutically targeted for HIV, IBD, and colorectal cancer therapy. Based on our findings, phytocannabinoids may offer a safer and more effective therapeutic option to modulate microbe-host epigenetic interactions in people with IBD and HIV.

Increased *NLRP6* expression in VEH/SIV RMs could be activated from the basolateral side by dsRNA intermediates formed during the HIV replication cycle along with LTA from dysbiotic gut bacteria. Such activation is strongly supported by the significantly high mRNA expression of *DHX15*, an RNA helicase that cooperates with *NLRP6* to recognize dsRNA and subsequently activate ISG expression through the mitochondrial antiviral signaling protein to exert antiviral effects. Promoter DNA hypermethylation coupled with significantly high *CYLD* mRNA expression in the CE of THC/SIV RMs suggests that phytocannabinoids can act upstream at the level of the DNA to reduce *NLRP6* gene/protein expression and downstream by upregulating *CYLD* to deubiquitylate and reduce NLRP6 protein activity. Further evidence supporting THC’s ability to inhibit NLRP6 activation comes from our recently published studies ^39,44^ describing significantly reduced mRNA expression of several ISGs and defensins, antimicrobial peptides produced by Paneth cells under the regulatory control of *NLRP6*.

Although NLRP6 has been described to activate both pyroptosis and necroptosis ^50^, the significantly high mRNA expression of *RIPK1, RIPK3, MLKL, PPM1B, TRPM7, ADAM17,* and *HMGB1* suggested that necroptosis is likely the dominant cell death pathway potentially activated by NLRP6 in the CE. Further, significantly high protein expression of p-RIPK3(Ser199) and p-MLKL(Thr357/Ser358) together with increased protein expression of HMGB1, provided strong evidence for the activation of necroptosis. More strikingly, excessive activation of proteins associated with the necroptosis pathway was detected in the intestine even when SIV replication was effectively suppressed by cART. As the terminal and crucial mediator of necroptosis, the significantly high expression of p-MLKL(Thr357/Ser358) to our knowledge, for the first time, identifies necroptosis as an important mechanism that could potentially drive intestinal epithelial barrier disruption in PLWH on suppressive cART. Furthermore, the presence of high HMGB1 protein expression means that necroptotic cell death could release HMGB1 into the extracellular space thereby activating and further amplifying secondary inflammatory responses leading to more widespread epithelial damage. Most importantly, the reduced mRNA and protein expression of necroptosis-associated proteins in the intestinal epithelium of cART-naïve and experienced THC/SIV RMs may be a direct consequence of downregulating NLRP6 (potentially other inflammasome sensors too) protein expression via promoter hypermethylation (upstream) and its functional activity by promoting its deubiquitylation through upregulating *CYLD* (downstream) (Figure 7). Given that the MB21D1 or cGAS-STING pathway also activates necroptosis^56^, THC-induced hypermethylation of MB21D1 provides an additional layer of protection against necroptosis.

Despite these clinically relevant findings, our study does have limitations. First, statistically significant NLRP6 upregulation was not detected in the colon of cART-naïve RMs. This may be attributed to the variation associated with between-group cross-sectional sampling performed on multiple animals at a single time point. However, in the cART-treated cohort, the collection of surgical intestinal resections from the same animals before and after SIV infection allowed the longitudinal detection of increased NLRP6 protein expression during acute infection that continued to remain high into the chronic stage despite viral suppression by cART. Our data also confirmed stronger NLRP6 expression in the jejunum than the colon and therefore should receive more scrutiny in the future. From a functional perspective, persistent NLRP6 and necroptosis activation in the jejunum could be more detrimental as it could interfere with nutrient and cART drug absorption, leading to incomplete viral suppression, increased systemic inflammation, and malnutrition. Moreover, the use of bulk CE made it impossible to tease apart the specific enterocyte populations that were most impacted by the methylation changes. Our future studies will utilize single-cell multiomics to investigate cell type-specific differences in gene expression and chromatin accessibility changes within the same cell population.

In conclusion, our findings provide an unprecedented insight into the epigenetic mechanisms regulating aberrant CE gene expression in chronic HIV/SIV infection, particularly in the paCGI of CE genes during chronic HIV/SIV infection, that impact inflammatory responses, cellular adhesions, and CE proliferation, including gene expression changes in the machinery responsible for these methylation changes. Due to the heritable nature of epigenetic marks, we postulate that these changes might partly explain a potential mechanism that drives chronic intestinal inflammation and barrier dysfunction in PLWH. Given the absence of FDA-approved inhibitors of NLRP6, we propose phytocannabinoids such as low-dose THC as a safe, feasible, inexpensive, and effective strategy to modulate inflammasome formation via epigenetic alteration of inflammasome sensors (i.e., *NLRP6*). Accordingly, modulation of inflammasome activation and downstream inhibition of necroptosis in the intestine as a therapeutic modality will benefit not only PLWH but also those suffering from other chronic inflammatory diseases like IBD ^73,74^. The important finding that CBD also blocked NLRP6 protein upregulation in vitro indicates its potential use as an alternative to THC and presents the opportunity to be combined with THC in appropriate ratios to reduce the psychotropic effects of THC while blocking NLRP6 hyperactivation. It is imperative that the hypermethylation of *NLRP6* and *MB21D1* not be deemed too detrimental to intestinal homeostasis, as such a therapeutic approach may be needed to suppress persistent intestinal inflammation in people with HIV and IBD. Nevertheless, given the reversible nature of epigenetic changes, discontinuation of phytocannabinoids alone after inflammation resolution should lead to restoration of normal methylation levels.

## Materials and Methods

### Animal care, ethics, and experimental procedures

All experiments using rhesus macaques were approved by the Tulane and LSUHSC Institutional Animal Care and Use Committee (Protocols 3574, 3581, and 3781). The Tulane National Primate Research Center (TNPRC) is an association for Assessment and Accreditation of Laboratory Animal Care International-accredited facility (AAALAC #000594). The NIH Office of Laboratory Animal Welfare assurance number for the TNPRC is A3071-01. All clinical procedures, including administration of anesthesia and analgesics, were carried out under the direction of a laboratory animal veterinarian. Animals were pre-anesthetized with ketamine hydrochloride, acepromazine, and glycopyrrolate, intubated and maintained on a mixture of isoflurane and oxygen. All possible measures were taken to minimize the discomfort of all the animals used in this study. Tulane University complies with NIH policy on animal welfare, the Animal Welfare Act, and all other applicable federal, state and local laws.

### Animal model and experimental design

Fifty-eight age and weight-matched male Indian RMs were randomly distributed into six groups. Group 1 [uninfected controls (n = 16)] served as uninfected controls. Groups 2-5 (n = 42) were infected intravenously with 100 times the 50% tissue culture infective dose (100TCID_50_) of SIVmac251. Groups 2 [VEH/SIV] (n=14) and 4 [VEH/SIV/ART] (n=8) received twice daily injections of vehicle (VEH) (1:1:18 of emulphor:ethanol:saline), intramuscularly. Groups 3 [THC/SIV] (n=12) and 5 [THC/SIV/ART] (n=8) received twice daily injection of THC intramuscularly four (THC/SIV) or two weeks (THC/SIV/ART) before SIV infection at 0.18mg/kg, as used in previous studies ^8^. This dose of THC was increased to 0.32mg/kg over a period of 2 weeks and maintained for the duration of the study. For animals in groups 4 and 5 (n = 16), cART was given daily by subcutaneous injection and included PMPA (20mg/kg) (Gilead), Emtricitabine (30mg/kg) (Gilead) and Dolutegravir (3mg/kg) (ViiV Healthcare) and was initiated two weeks post-infection. As mentioned previously^44^, four macaques in group 4 and four macaques in group 5 received two injections of anti-α4β7 integrin (LN60, LC48, LM85, LJ21) or control IgG (LM56, LH75, LH92, LI81) (50 mg/kg of anti-α4β7 or control IgG) beginning 4 months post infection (MPI) at three-week intervals as part of another study before the jejunum resections were collected at 5 MPI.

Colon tissue (∼5cm) was collected for epithelial cell isolation at necropsy from all animals except animals from groups 4 and 5. For these animals, jejunum resections (5∼cm) were collected at pre-infection, 1 MPI and 5 MPI. For histopathological and immunohistochemical evaluation, colon and jejunum tissues were fixed in Z-fix, embedded in paraffin, sectioned at 5 µM for further analysis.

SIV levels in plasma, colon, and jejunum were quantified using the TaqMan One-Step Real-time RT-qPCR assay that targeted the LTR gene ^75^. SIV inoculum, infection duration, plasma/colon/jejunum viral loads, and colon/jejunum histopathology can be found in Table 1.

### Colonic epithelial cell isolation and DNA/RNA extraction

Colon epithelial cells were isolated ^38^ for Arraystar microarray ^44^ and RRBS. The purified epithelial components were then collected in RNAlater (Thermo Fisher Scientific) for DNA extraction (DNeasy Blood and Tissue Kit; Qiagen Inc, CA), or lyzed in Qiazol (Qiagen Inc, CA) for total RNA extraction.

### Reduced representation bisulfite sequencing (RRBS) library construction

Methyl-MiniSeq Library preparation was performed by Zymo Research. Libraries were prepared from 200-500ng of genomic DNA digested with 60 units of Taqαl and 30 units of MspI (NEB) sequentially and then extracted with DNA Clean & Concentrator^TM^-5 (Zymo Research). Fragments were ligated to pre-annealed adapters containing 5’-methyl-cytosine according to Illumina’s specified guidelines (www.illumina.com). Adapter-ligated fragments of 150-250bp and 250-300bp were recovered from a 2/5% NuSieve 1:1 agarose gel (Zymoclean Gel DNA Recovery Kit, Zymo Research). The fragments were then bisulfite treated using EX DNA Methylation-Lightning Kit (Zymo Research). Preparative-scale PCR was performed, and the resulting products purified for sequencing on an Illumina HiSeq.

Sequence reads from bisulfite-treated EpiQuest libraries were identified using standard Illumina based-calling software and then analyzed using Zymo Research propietrary analysis pipeline, which is written in Python along with Bismark (www.bioinfomatics.babraham.ac.uk/projects/bismark/) to perform the alignment. Index files were constructed using the *bismark_genome_preparation*) command and the entire reference genome. The *non-directional* parameter was applied while running Bismark. All other parameters were set to default. Filled-in nucleotides were trimmed off while doing methylation calling. The methylation level of each sampled cystosine (C) was estimated as the number of reads reporting a C, divided by the total number of reads reporting a C or thymine. Fisher’s exact test or *t* test was performed for each CpG site which has at least five reads coverage, and promoter, gene body, and CpG island annotations were added for each CpG included in the comparison.

### Global mRNA profiling

Sample preparation, microarray analysis, and statistics were performed by Arraystar, as previously described ^44^.

### GO enrichment analysis

To determine the biological functions of differentially methylated genes, Gene Ontology (GO) enrichment analysis was carried out as previously decreased (52), but *Macaca mulatta* was the selected organism.

### Quantitative image analysis of colon and jejunum sections

Colon and jejunum sections from animals in Table 1 were stained with antibodies specific for NLRP6 (Millipore Sigma: ABF29; colon 1:100 dilution), p-RIPK3(Ser199) (MyBiosource: MBS1569089; 1:200 dilution), PPM1B (Proteintech: 67647-1-Ig; 1:500 dilution), p-MLKL(Thr357) (Novus Biologicals: MAB91871, 1:200 dilution), p-MLKL(Ser358) (GeneTex; GTX00973, 1:100 dilution), HMGB1 (D3E5) (Cell Signaling Technology: 6893; 1:50 dilution). At least eight areas per tissues for each RM was scanned using a Zeiss LSM800 confocal microscope (Carl ZEISS Microscopy, LLC) at 20X objective, and imported as digital images into HALO software (Indica Labs) for image quantitation analysis. Since cells in the lamina propria also stained positive for all protein markers, regions of interest (ROIs) were manually drawn around the colon or jejunum epithelium using the HALO pen tool. The Indica Labs’ Area quantification module (FL v4.2.3) was then utilized to determine the average intensity of NLRP6 in each epithelial cell, in the ROIs. The output values (average positive area fluorescence) were used to calculate the total fluorescence within the ROIs for colonic and jejunal sections.

## Data availability

RRBS data will be submitted to Gene Expression Omnibus (GEO), and accession numbers will be provided if we are approved for resubmission. mRNA profiling data was previously submitted to GEO (accession no: GSE223482; https://www.ncbi.nlm.nih.gov/geo/query/acc.cgi?acc=GSE223482) ^44^.

## Data analysis

Distribution of differentially methylated site in the paCGI region of protein-coding genes and the average methylation change that occurred within NLRP6 paCGI, mRNA gene expression changes, and NLRP6 and necroptosis-associated protein image quantitation in the colonic and jejunum epithelium and lamina propria mononuclear cells were graphed and analyzed using Prism v9 software (GraphPad).

After verifying data assumptions (normal distribution) using Anderson-Darling, D’Agostino & Pearson, Shapiro-Wilk, and Kolmogorov-Smirnov tests, p-values were calculated using either Kruskal-Wallis (differential methylation change in paCGIs and NLRP6 fluorescence in colonic LPL) or Ordinary one-way ANOVA (Area fluorescence of NLRP6, p-RIPK3(Ser199), PPM1B, p-MLKL(Thr357), p-MLKL(Ser358) and HMGB1 in colon and jejunum epithelium), unpaired *t* test (VEH/SIV relative to control and THC/SIV RMs methylation ratio of DMSs within NLRP6 and MB21D1 paCGI), Mann-Whitney U test (THC/SIV relative to control RMs methylation ratio of DMSs within NLRP6 paCGI), or Mixed-effects analysis (NLRP6 area fluorescence in jejunum epithelial cells. Heatmaps were made with TB Tools ^76^.

## Supporting information

Supplemental Table 1 and Supplemental Figures 1, 2 and 3 to be used for link to the file on the preprint site

## Acknowledgements

The authors would like to thank Ronald S. Veazey, Maurice Duplantis, Faith R. Schiro, Cecily C. Midkiff, Coty Tatum (Tulane National Primate Research Center, Covington, Louisiana) and Eunhee Lee (Texas Biomedical Research Institute, San Antonio, Texas) for their technical assistance with nonhuman primate studies.

## Author Contributions

LP and MMW equally contributed to the overall gene expression and DNA methylation data processing, analysis, immunofluorescence studies, image analysis, cell culture and writing of the manuscript. MM carried out the overall planning, direction, and design of the in vivo and in vitro experiments. LP, MMW, LR, BG, JAD and MM carried out the day-day sampling scheduling (animal experiments) and prepared the samples for RRBS and microarray gene expression, and performed immunofluorescence/image quantification, and data analysis. BL and CMO assisted with viral load assays, data analysis, and overall data interpretation and conclusions. LP, MMW and MM wrote the manuscript with input from all authors. JAD, BL, and CMO provided helpful suggestions and review of the manuscript. All authors read and approved the final version of the manuscript.

---

**Supplemental Table S1.** Number of CpG sites hypomethylated (in red) or hypermethylated (in black) within CpG islands of unique genes in VEH/SIV and THC/SIV, compared to controls and THC/SIV rhesus macaques.

**Supplemental Figure S1.** Flow chart showing stepwise processing of RRBS DNA methylation data and identification of differentially methylated sites in paCGIs.

**Supplemental Figure S2.** Clustering heatmaps of the top 100 differentially methylated CpG sites in VEH/SIV (**A**) and THC/SIV (**B**) relative to control rhesus macaques (RMs) and in VEH/SIV (**C**) compared to THC/SIV RMs. Pearson’s correlation coefficients for all CpG sites between VEH/SIV vs control (**D**), THC/SIV vs control (**E**), and VEH/SIV vs THC/SIV (**F**) RMs were 0.9400, 0.9358, and 0.9359, respectively.

**Supplemental Figure S3.** UCSC Genome track for *MB21D1* showing the location of the 108 bp CGI (**A**). p-RIPK3(Ser199) (green) (**B-D**), and PPM1B (green) (**F-H**) protein expression in jejunum tissues before SIV infection (**B&F)**) and at 5 months post SIV infection in cART suppressed VEH/SIV (**C&G**) and THC/SIV (**D&H**) rhesus macaques. RIPK3(Ser199) (**E**) and PPM1B (**I**) protein quantification in jejunum. Immunofluorescence data were analyzed using one-way ANOVA followed by Tukey’s multiple comparison post hoc test. A p-value of <0.05 was considered significant. *TLR3* (**J**) gene expression in significantly upregulated in the colonic epithelium of VEH/SIV but not in THC/SIV compared to control rhesus macaques.

