## Supplemental Table 1 and Supplemental Figures 1, 2 and 3 to be used for link to the file on the preprint site for "Epigenetic modulation of NLRP6 inflammasome sensor as a therapeutic modality to reduce necroptosis-driven gastrointestinal mucosal dysfunction in HIV/SIV infection"

**Table S1:** Number of CpG sites hypomethylated (in red) or hypermethylated (in black) within CpG islands of unique genes in VEH/SIV and THC/SIV, compared to controls and THC/SIV RMs.

| Function | Gene | VEH-SIV vs Control | Average Methylation Difference | THC-SIV vs Control | Average Methylation Difference | VEH-SIV vs THC-SIV | Average Methylation Difference | Total number of CpG sites |
|---|---|---|---|---|---|---|---|---|
| Inflammatory response | CXCL12 | 5 | -21.2 | 5 | -23.8 | 1 | -16 | 257 |
| | TOLLIP | 3 | -40.7 | 2 | -48 | 1 | -11 | 130 |
| | NLRP6 | 8(6) | -6.6 | 6(38) | 12.5 | 12(1) | -18 | 248 |
| | LEP | 2(3) | 2.6 | 1(10) | 15.6 | 9 | -16 | 61 |
| | MB21D1 | 2 | -16.5 | 1(9) | 24.2 | 6 | -22 | 108 |
| Cellular adhesion | CLDN6 | 1(2) | 10.7 | (4) | 23 | 5 | -15.8 | 48 |
| | CLDN11 | 4 | -29.8 | 1 | -29 | 2 | -12 | 139 |
| | ICAM1 | 5 | -18.2 | 3 | -20 | 2 | -15 | 132 |
| | PRKCB | (1) | 11 | (8) | 17.4 | 3 | -18.3 | 134 |
| | PCDH17 | 3(9) | 3.7 | 1(52) | 24.1 | 29 | -19.3 | 229 |
| | PCDH19 | 2 | -34.5 | 2(1) | -8.33 | 8(1) | -22.6 | 368 |
| | ADRB1 | 1(5) | 12.3 | (12) | 29 | 7 | -21 | 238 |
| | CDH7 | 2(2) | -3 | 1(20) | 24.9 | 17 | -22.9 | 108 |
| Oxidative stress | CYGB | 2(1) | -10 | 2(3) | -1.6 | 6 | -12.7 | 27 |
| | GPX3 | 2 | -17.5 | 2 | -15.5 | 1 | -11 | 40 |
| Colonic epithelial cell proliferation | WIF1 | 1(1) | -20.5 | 1(7) | 8.25 | 3 | -22 | 91 |
| | SFRP1 | 3(1) | -11.5 | 3(18) | 11.3 | 12 | -16.9 | 131 |
| | TERT | 4(1) | -19.5 | 3(10) | 18.6 | 7 | -28 | 243 |
| | HAND2 | 1 | -11 | (1) | 16 | 4 | -16.5 | 131 |
| Apoptosis | DAPK1 | 4 | -27.8 | 4(1) | -16 | 6 | -21.8 | 145 |
| | BCL2L1 | 1 | -19 | 1 | -19 | -- | -- | 118 |

**Figure S1**

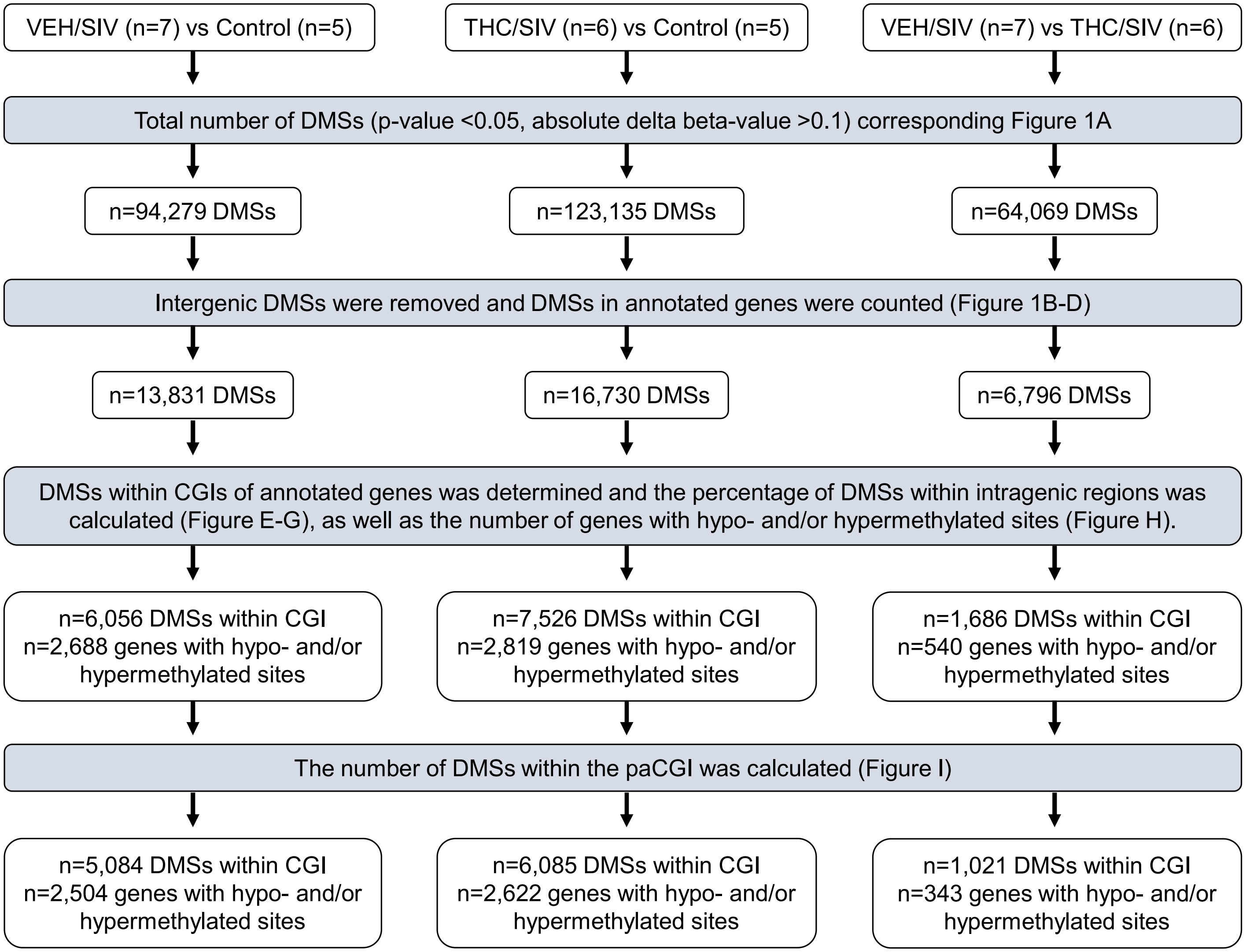
VEH/SIV (n=7) vs Control (n=5)
THC/SIV (n=6) vs Control (n=5)
VEH/SIV (n=7) vs THC/SIV (n=6)
Total number of DMSs (p-value <0.05, absolute delta beta-value >0.1) corresponding Figure 1A
n=94,279 DMSs
n=123,135 DMSs
n=64,069 DMSs
Intergenic DMSs were removed and DMSs in annotated genes were counted (Figure 1B-D)
n=13,831 DMSs
n=16,730 DMSs
n=6,796 DMSs
DMSs within CGIs of annotated genes was determined and the percentage of DMSs within intragenic regions was calculated (Figure E-G), as well as the number of genes with hypo- and/or hypermethylated sites (Figure H).
n=6,056 DMSs within CGI n=2,688 genes with hypo- and/or hypermethylated sites
n=7,526 DMSs within CGI n=2,819 genes with hypo- and/or hypermethylated sites
n=1,686 DMSs within CGI n=540 genes with hypo- and/or hypermethylated sites
The number of DMSs within the paCGI was calculated (Figure I)
n=5,084 DMSs within CGI n=2,504 genes with hypo- and/or hypermethylated sites
n=6,085 DMSs within CGI n=2,622 genes with hypo- and/or hypermethylated sites
n=1,021 DMSs within CGI n=343 genes with hypo- and/or hypermethylated sites

Figure S2

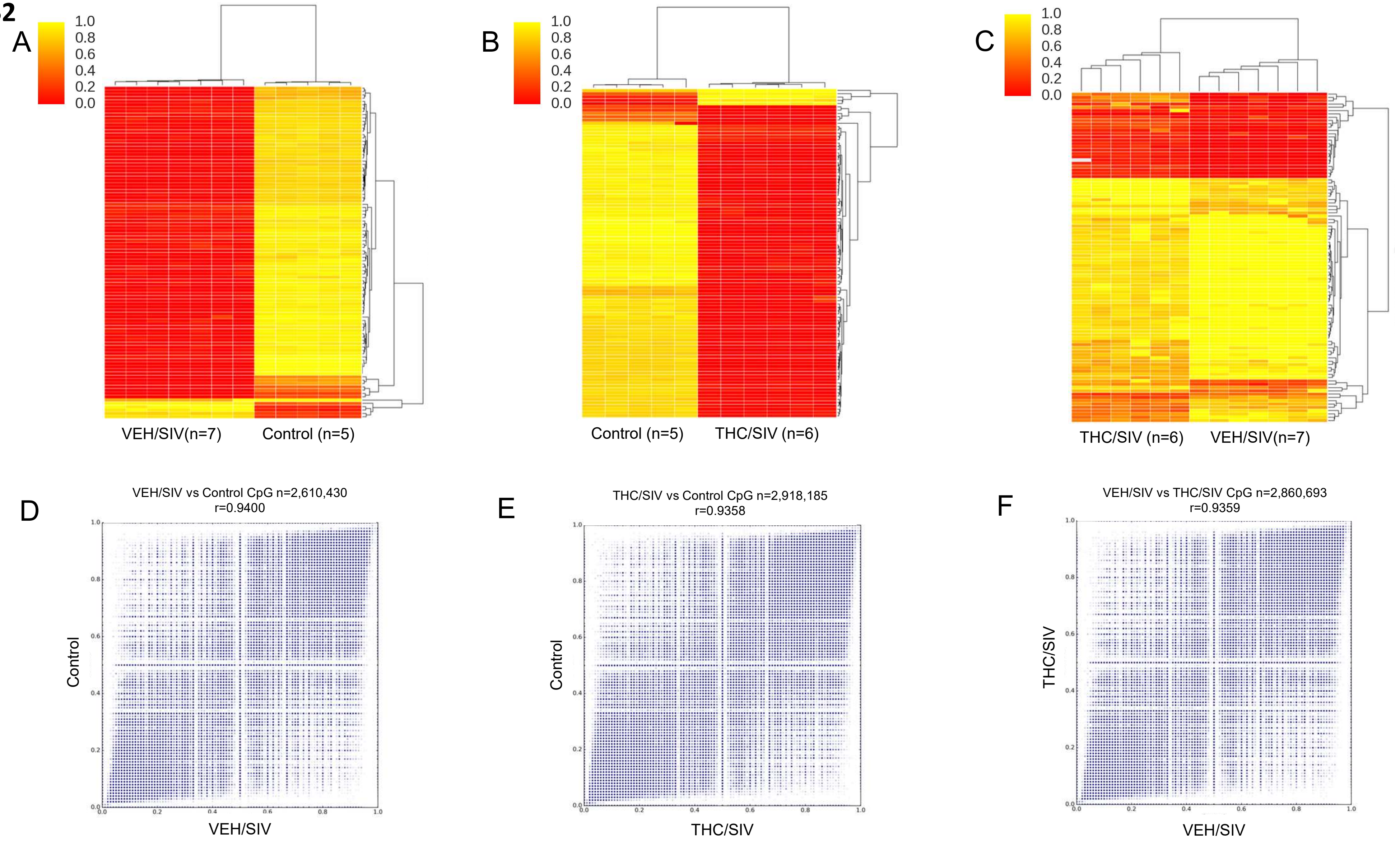
A
B
C
1.0
0.8
0.6
0.4
0.2
0.0
VEH/SIV(n=7)
Control (n=5)
Control (n=5)
THC/SIV (n=6)
THC/SIV (n=6)
VEH/SIV(n=7)
D
VEH/SIV vs Control CpG n=2,610,430
r=0.9400
Control
VEH/SIV
E
THC/SIV vs Control CpG n=2,918,185
r=0.9358
Control
THC/SIV
F
VEH/SIV vs THC/SIV CpG n=2,860,693
r=0.9359
THC/SIV
VEH/SIV

### Figure S3

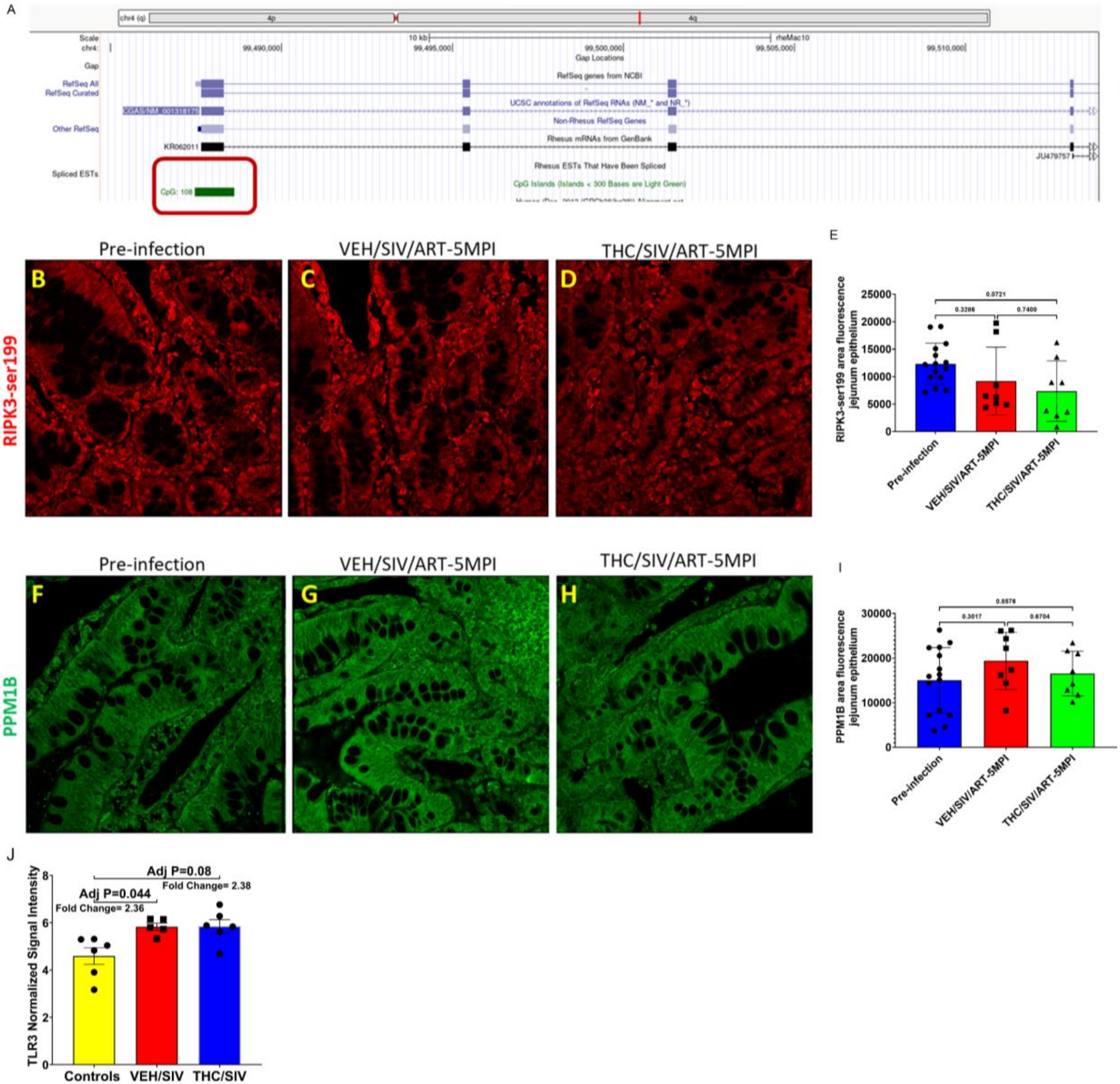

A
chr4 (q)
4p
4q
Scale
chr4:
10 kb
rheMac10
99,490,000
99,495,000
99,500,000
99,505,000
99,510,000
Gap
Gap Locations
RefSeq All
RefSeq Curated
RefSeq genes from NCBI
UCSC annotations of RefSeq RNAs (NM_* and NR_*)
Other RefSeq
Non-Rhesus RefSeq Genes
KR062011
Rhesus mRNAs from GenBank
JU479757
Spliced ESTs
Rhesus ESTs That Have Been Spliced
CpG: 108
CpG Islands (Islands < 300 Bases are Light Green)
Pre-infection
VEH/SIV/ART-5MPI
THC/SIV/ART-5MPI
B
C
D
RIPK3-ser199
E
RIPK3-ser199 area fluorescence jejunum epithelium
25000
20000
15000
10000
5000
0
0.0721
0.3286
0.7400
Pre-infection
VEH/SIV/ART-5MPI
THC/SIV/ART-5MPI
Pre-infection
VEH/SIV/ART-5MPI
THC/SIV/ART-5MPI
F
G
H
PPM1B
I
PPM1B area fluorescence jejunum epithelium
30000
20000
10000
0
0.8576
0.3017
0.6704
Pre-infection
VEH/SIV/ART-5MPI
THC/SIV/ART-5MPI
J
TLR3 Normalized Signal Intensity
8
6
4
2
0
Adj P=0.08
Fold Change= 2.38
Adj P=0.044
Fold Change= 2.36
Controls
VEH/SIV
THC/SIV
